# The integrated stress response mediates type I interferon driven necrosis in *Mycobacterium tuberculosis* granulomas

**DOI:** 10.1101/499467

**Authors:** Bidisha Bhattacharya, Shiqi Xiao, Sujoy Chatterjee, Michael Urbanowski, Alvaro Ordonez, Elizabeth A. Ihms, Garima Agrahari, Shichun Lun, Robert Berland, Alexander Pichugin, Yuanwei Gao, John Connor, Alexander Ivanov, Bo-Shiun Yan, Lester Kobzik, Sanjay Jain, William Bishai, Igor Kramnik

**Author notes:** Johns Hopkins School of Medicine (Shiqi Xiao, Michael Urbanowski, Alvaro Ordonez, Elizabeth A. Ihms, Shichun Lun, William Bishai). Contributed equally (Bidisha Bhattacharya, Shiqi Xiao).

## Abstract

Necrosis in the tuberculous granuloma is a hallmark of tuberculosis that enables pathogen survival and transmission. Susceptibility to tuberculosis and several other intracellular bacteria is controlled by a mouse genetic locus, *sst1*, and mice carrying the sst1-suscepible (sst1S) genotype develop necrotic inflammatory lung lesions, similar to human TB granulomas. Our previous work established that increased disease severity in sst1S mice reflects dysfunctional macrophage effector or tolerance mechanisms, but the molecular mechanisms have remained unclear. Here we demonstrate that sst1S macrophages develop aberrant, biphasic responses to TNF characterized by super-induction of stress and type I interferon pathways after prolonged TNF stimulation with this late-stage response being initiated by oxidative stress and Myc activation and driven via a JNK - IFNβ - PKR circuit. This circuit leads to induction of the integrated stress response (ISR) mediated by eIF2α phosphorylation and the subsequent hyper-induction of ATF3 and ISR-target genes Chac1, Trib3, Ddit4. The administration of ISRIB, a small molecule inhibitor of the ISR, blocked the development of necrosis in lung granulomas of *M. tuberculosis-infected* sst1S mice and concomitantly reduced the bacterial burden revealing that induction of the ISR and the locked-in state of escalating stress driven by type I IFN pathway in sst1S macrophages plays a causal role in the development of necrosis. Our data support a generalizable paradigm in intracellular pathogen-host interactions wherein host susceptibility emerges within inflammatory tissue due to imbalanced macrophage responses to growth, differentiation, activation and stress stimuli. Successful pathogens such as *M. tuberculosis* may exploit this aberrant response in susceptible hosts to induce necrotic lesions that favor long-term pathogen survival and transmission. Interruption of the aberrant stress response with inhibitors such as ISRIB may offer novel therapeutic strategies.

## INTRODUCTION

Susceptibility to infectious agents varies considerably within populations of natural hosts, and manifest as a spectrum of severity and outcomes of infectious diseases. These heterogeneous outcomes are governed by multi-dimensional interactions of genetic, behavioral and environmental factors. A corollary is that key pathogenic properties of microbes are best revealed within the specific context of susceptible hosts. Ultimately, understanding general and specific mechanisms of host susceptibility to individual pathogens or groups of pathogens is vital for the development of effective preventive and therapeutic strategies, including against infectious diseases caused by antibiotic-resistant bacteria.

As a product of co-evolution with their natural hosts, the highly specialized human pathogen *Mycobacterium tuberculosis* (Mtb) has acquired the ability to create local environments within susceptible hosts conducive for pathogen survival, replication and transmission. It targets the lungs of immunocompetent hosts to form *de novo* structures – granulomas – that initially provide a sanctuary for the pathogen and, at the same time, protect the host from disseminated and potentially lethal infection^1,2^. Ultimately, TB progression leads to massive destruction of lung TB granulomas and to formation of lung cavities that are essential for the pathogen transmission via aerosols – the only epidemiologically significant route of transmission within human populations.

So far, susceptibility to Mtb infection has been largely viewed as an innate or induced defect of host immunity^3^. Less understood, are intrinsic mechanisms of susceptibility that develop within lung tissue in otherwise immunocompetent individuals. Indeed, in TB granulomas, the acquisition of permissiveness to Mtb infection by certain host cells does not equate to loss of control mechanisms at the whole organism level^4^.

Host mechanisms that control tissue damage are collectively referred to as tolerance to infection, which is defined at the whole organism level as host ability to maintain a certain health status in the presence of substantial pathogenic loads^5,6^. At the tissue level, tolerance is determined by cell resilience to stressors within inflammatory milieu induced by pathogens’ invasion^7^. Indeed, mechanisms essential for host resistance to many bacterial infections, such as reactive oxygen and nitrogen species, may also induce collateral tissue damage, unless they are countered by host mechanisms of stress resilience. The salient feature of host tolerance strategies, as opposed to resistance, is their focus on maintaining host cell and tissue homeostasis^8,9^, and, therefore, tolerance mechanisms are expected to operate in pathogenesis-specific rather than pathogen-specific manner.

To study mechanisms underlying host susceptibility to pulmonary TB, we developed a mouse model of human-like necrotic TB granulomas using C3HeB/FeJ (C3H) mice^10–12^. Using forward genetic analysis in a cross of the C3H with relatively TB resistant C57BL/6J (B6) mice, we have identified a novel genetic locus *sst1* (<u>***s***</u>uper-<u>***s***</u>usceptibility to <u>***t***</u>uberculosis <u>***1***</u>), as a specific determinant of necrosis in TB granulomas^13^. Congenic mice that carry the C3HeB/FeJ-derived *sst1* susceptibility allele (sst1S) on the resistant B6 background, B6.*Sst1^C3H,S^* (abbreviated B6-sst1S), developed large, well-organized pulmonary necrotic granulomas after a low dose aerosol Mtb infection, even though these mice initially controlled Mtb replication similarly to the parental B6 mice^14^. The necrosis in the B6-sst1S TB lesions occurred with bacterial loads approximately 50-fold lower than in the parental C3HeB/FeJ mice indicating that extreme bacterial loads did not drive the sst1S-mediated necrosis. These data demonstrated for the first time that mechanisms controlling Mtb-inflicted necrotic damage could be genetically uncoupled from the effector immune mechanisms controlling the bacterial load.

Subsequent experiments using the *sst1* congenic mice revealed that the *sst1* locus controlled macrophage interactions with Mtb and other intracellular bacterial pathogens – *Listeria monocytogenes*^15^ and *Chlamydia pneumonia^16^.* Using positional cloning approach, we have identified a variant of the interferon-inducible nuclear protein Sp110 or Ipr1 (intracellular pathogen resistance 1), as a strong candidate gene, whose expression is completely abolished in sst1S macrophages^17^. The human homologue of Ipr1 – Sp110b – has been shown to play important roles in immunity to infections^18^ and in control of macrophage activation^19^. However, the molecular and cellular mechanisms explaining the role of *sst1/Sp110* in the pathogenesis of TB and other infections remain to be elucidated.

Here we present evidence that the mouse genetic locus *sst1* is a broad determinant of host tolerance as it controls macrophage resilience to stress induced by tumor necrosis factor (TNF). TNF is a major cytokine required for innate and adaptive immune responses to many infectious agents. It is especially important for the formation and maintenance of TB granulomas^20^. Compared to the wild type, TNF stimulation of the sst1S congenic macrophages induced an aberrant activation cascade beginning with elevated proteotoxic stress, Myc hyperactivity and super-induction of type I interferon (IFN-I), and culminating with an escalating integrated stress response (ISR) mediated by interferon-inducible eIF2α kinase – protein kinase R (PKR) (Supplemental Figure 1). This dysregulated macrophage response to TNF provoked susceptibility to two taxonomically unrelated intracellular bacterial pathogens, *Francisella tularensis* LVS and *Mycobacterium tuberculosis.* Thus, our findings delineate a common mechanism of TNF-induced macrophage damage and increased susceptibility to intracellular bacterial pathogens sculpted by hyperactivity of type I IFN and sustained activation of the integrated stress response pathway.

## RESULTS

### TNF triggers hyperactivity of type I interferon (IFN-I) and stress pathways in sst1S macrophages

Previously we reported that B6-sst1S mice infected with *Chlamydia pneumoniae*, as well as their bone marrow-derived macrophages (BMDMs) infected with *C.pneumoniae*, produce higher levels of IFNβ^16^. This was associated with increased death of infected macrophages in vitro, which could be reduced using IFN receptor (IFNAR1) blocking antibodies. To dissect mechanisms behind the upregulated IFNβ production, we compared IFNβ secretion by the B6wt and B6-sst1S BMDMs, stimulated either with TNF (which induces low levels of IFNβ in B6 macrophages^21^) or a classical IFNβ inducer poly(I:C). The B6-sst1S macrophages secreted higher levels of IFNβ protein in response to both stimuli (**Fig 1A and Suppl. Fig. 2A**, respectively). Next, we compared the kinetics of TNF-induced IFNβ mRNA expression in B6wt vs B6-sst1S BMDMs. Initially, TNF induced similarly low levels of IFNβ mRNA expression in both cell types. Then, while IFNβ levels remained relatively stable in B6wt macrophages, in the B6-sst1S cells the IFNβ mRNA expression significantly increased between 8 – 24 h, such that the strain difference in IFNβ mRNA levels reached 10-20-fold by 24 h (**Fig. 1B**). In addition, the B6-sst1S macrophages stimulated with TNF expressed significantly higher levels of the interferon-stimulated gene Rsad2 (viperin) (**Suppl. Fig.2B**), whose up-regulation was significantly reduced (by 70-75%) in the presence of IFNAR1 blocking antibodies, thus, confirming activation of the type I IFN (IFN-I) pathway in the B6-sst1S cells. The IFNβ and Rsad2 mRNA expression kinetics demonstrated that the bias towards IFN-I pathway activation in the B6-sst1S macrophages occurred at a later stage of persistent stimulation with TNF.

**Figure 1.**
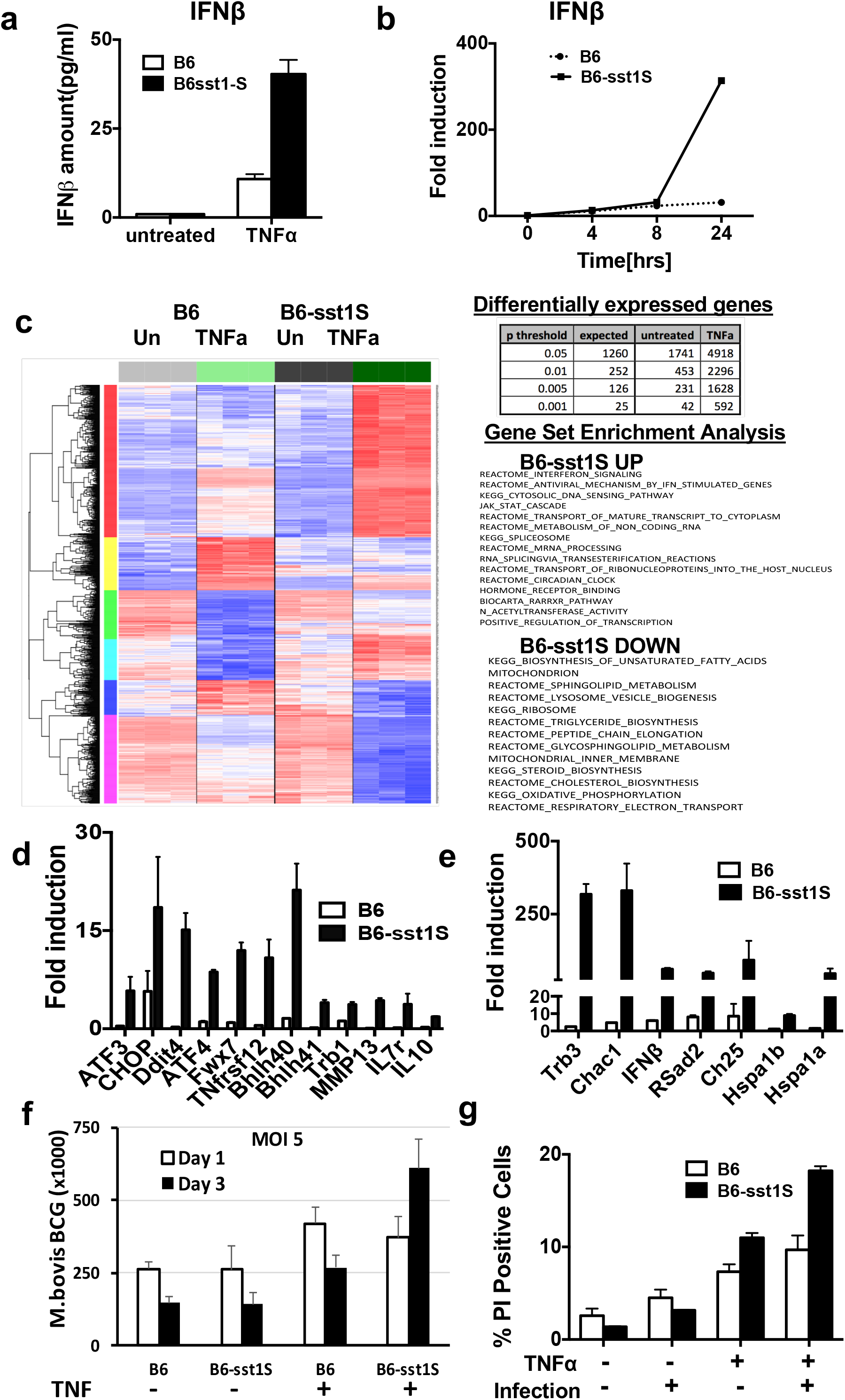
Super-induction of IFNβ in B6-sst1S BMDM after prolonged stimulation with TNF. **a)** IFNβ protein concentration in supernatants of B6wt and B6-sst1S BMDM treated with l0ng/ml TNFa for 24 h was detected using ELISA. Results represent data from two independent experiments. **b)** Timecourse of IFNβ mRNA expression in B6-sst1S and B6wt BMDMs after treatment with10ng/mL, as determined using real time qRT-PCR. The data are representative of three biological replicas. **c)** Comparison of gene expression profiles of B6-sst1S vs B6 BMDMs stimulated with TNF (10ng/mL) for 18 h using hierarchical clustering and gene set expression analysis (GSEA). The global gene expression was determined using Affimetrix GeneChip Mouse Gene 2.0 Arrays. **d)** Validation of microarray data using gene-specific real time qRT-PCR. **e)** Differential stress response and IFN-I pathways gene expression in TNF-stimulated B6-sst1S and B6wt BMDMs. **f)** Effect of TNF priming on bacterial loads in B6-sst1S and B6 BMDMs 1 and 3 days after infection with *M.bovis* BCG. **g)** The sst1-dependent effect of TNF priming on macrophage survival 24 h after infection with *F. tularensis* LVS. The B6-sst1S and B6wt BMDMs were pre-treated with TNF (10 ng/ml) for 18 h and infected with *F.t.* LVS at MOI =1. Cell death was measured using *%* of PI positive cells.

To characterize effects of the *sst1* locus on the late response of primary macrophages to TNF more broadly we compared transcriptomes of B6 and B6-sst1S BMDMs after 18 h of stimulation with TNF at 10 ng/ml (**Fig. 1C**). While no significant differences were detected in naive macrophages, the gene expression profiles of TNF-treated cells diverged substantially with 492 genes differentially expressed at p<0.001 (**Table in Fig.1C**). The most prominent differentially expressed cluster was composed of genes that were selectively upregulated by TNF in B6-sst1S, but not B6wt macrophages (**Fig. 1C**). Using Gene Set Enrichment Analysis (GSEA) we found significant enrichment for the type I interferon-regulated genes in sst1S macrophages responding to TNF. Genes involved in nuclear RNA processing and nucleo-cytoplasmic transport were also upregulated by TNF in sst1S macrophages. Strikingly, multiple biosynthetic pathways were coordinately downregulated in TNF-stimulated sst1S macrophages, including lipid and cholesterol biosynthesis, protein translation, ribosome, mitochondrial function and oxidative phosphorylation. Further validation of the differential gene expression using quantitative real time RT-PCR (qRT-PCR), demonstrated up-regulation of a number of other pathogenically important genes, such as IL-10, Mmp-13, IL-7R, Death Receptor 3 (Dr3/Tnfrsf12), transcription factors Bhlh40 and Bhlh41 in the sst1S cells (**Fig. 1D**). Significant upregulation of IFNβ and typical IFN-I-inducible genes Rsad2 and Ch25h confirmed our previous observations of IFN-I hyperactivity. Also, a group of genes (Atf3, Chop10, Ddit4, Trib3 and Chac1) induced during integrated stress response (ISR), as well as markers of proteotoxic stress (PS) Hspa1a and Hspa1b, were significantly enriched among the upregulated genes, and their upregulation by TNF specifically in sst1S macrophages was confirmed using qRT-PCR (**Fig. 1E**).

To explore functional consequences of the aberrant activation of the sst1S macrophages by TNF, we infected them in vitro with attenuated intracellular bacteria *Mycobacterium bovis* BCG or *Francisella tularensis* LVS. Indeed, the ability to reduce the bacterial loads within 3 days of infection was specifically abrogated by TNF pre-treatment in the sst1S macrophages (**Fig. 1F**). Similarly, TNF pre-treatment increased macrophage susceptibility in the *F.tularensis* LVS infection model. Assessing the levels of IFNβ, Rsad2 and integrated stress response mRNAs Chac1 and Trb3 in either naïve of TNF-pretreated, we observed that TNF treatment made a more significant contribution to IFN and stress pathways induction, as compared to the bacteria alone (**Suppl. Fig. 2C**). The bacterial loads also increased in TNF pre-treated susceptible macrophages suggesting that stress compromised macrophage resistance to *F. tularensis* LVS as well (**Suppl. Fig. 2D**). Finally, TNF pre-treatment increased death of *F.tularensis* LVS-infected macrophages to a greater degree in the sst1S background (**Fig. 1G**).

To assess the impact of the *sst1* locus in vivo, we infected the *sst1* resistant and susceptible congenic mice via the respiratory route with 1600 CFU of *F. tularensis* LVS. The survival of *sst1S* mice was significantly lower compared to that of *sst1R* congenic mice (**Suppl. Fig.2E**). Importantly, the bacterial replication was initially similar in the lungs, spleens, and livers of both mouse strains. The bacterial loads, however, significantly diverged between days 5 and 11 reaching 100-fold higher levels in the organs of the *sst1S* mice (**Suppl. Fig. 3A**) which also displayed extensive necrotic lung inflammation by day 11 (**Suppl. Fig. 3B-C**). FACS analysis of inflammatory cells isolated from the infected lungs of immune competent mice eight days post infection demonstrated similar proportions of CD4+ and CD8+ T cell populations and NK cells. The proportions of activated CD69+ cells within those populations were also similar (**Table 1**). We observed substantial difference in the myeloid compartment (CD11b+), where the fraction of immature monocyte-like Ly6C+F4/80^-^ cells was significantly higher in the lungs of the *sst1S* mice (11.2%), as compared to 5.3% in the B6 lungs. The ratio of the more mature (Ly6C+F4/80+) monocyte-derived macrophages to the immature (Ly6C+F4/80^-^) cells in the lungs of the *sst1S* mice was 1.1, as compared to 3.7 in the resistant ones (**Table 2**). These data imply either delayed maturation of the newly recruited inflammatory monocytes in the lungs of the sst1S mice and/or more rapid turnover of monocyte-derived macrophages during *F. tularensis* LVS infection *in vivo*, which would be consistent with their more rapid demise, as observed in vitro. In addition, we detected a higher proportion of the IL-10 producing myeloid cells in the sst1S mouse lungs at that time, which is also in agreement with the sst1S BMDM phenotype observed in vitro (**Suppl. Fig. 3D**). Additional experiments were performed using *sst1* congenic *scid* mice developed in our laboratory. Both the sst1 R *scid* and sst1S *scid* mice succumbed to infections with 60 CFU of *F.t.* LVS. However, the survival time was shorter and the bacterial loads were 30-50-fold higher in the *sst1* susceptible *scid* mice (**Suppl. Fig.3E and 3F**) demonstrating that the innate immune defect was responsible for the sst1-mediated phenotype *in vivo.*

**Table 1.** Flow cytometry of lymphoid lung cells 8 days following aerosol challenge of C3H and C3H.B6-sst1 mice with 300 CFU of *F.t.* LVS

| Mouse strain | C3H |  | C3H.B6-sst1 |  |
| --- | --- | --- | --- | --- |
| Lymphocyte population | % of total | SD | % of total | SD |
| CD4<sup>+</sup> | 8.2 | 4.6 | 7.5 | 1.3 |
| % of CD69<sup>+</sup> | 25.7 | 9.2 | 28.1 | 6.9 |
| CD8<sup>+</sup> | 4.0 | 1.3 | 5.6 | 2.5 |
| % of CD69<sup>+</sup> | 36.2 | 7.2 | 33.7 | 5.5 |
| DX5<sup>+</sup>CD4<sup>-</sup> | 10.1 | 4.6 | 11.2 | 3.0 |
| % of CD69<sup>+</sup> | 7.7 | 1.6 | 7.4 | 0.6 |
| DX5<sup>+</sup>CD4<sup>+</sup> | 0.23 | 0.17 | 0.15 | 0.04 |
| % of CD69<sup>+</sup> | 26.2 | 2.5 | 30.2 | 6.5 |
| CD19<sup>+</sup> | 1.12 | 0.60 | 2.21 | 0.67 |
| % of I-A<sup>K+</sup> | 100 | 0 | 100 | 0 |

**Table 2.**
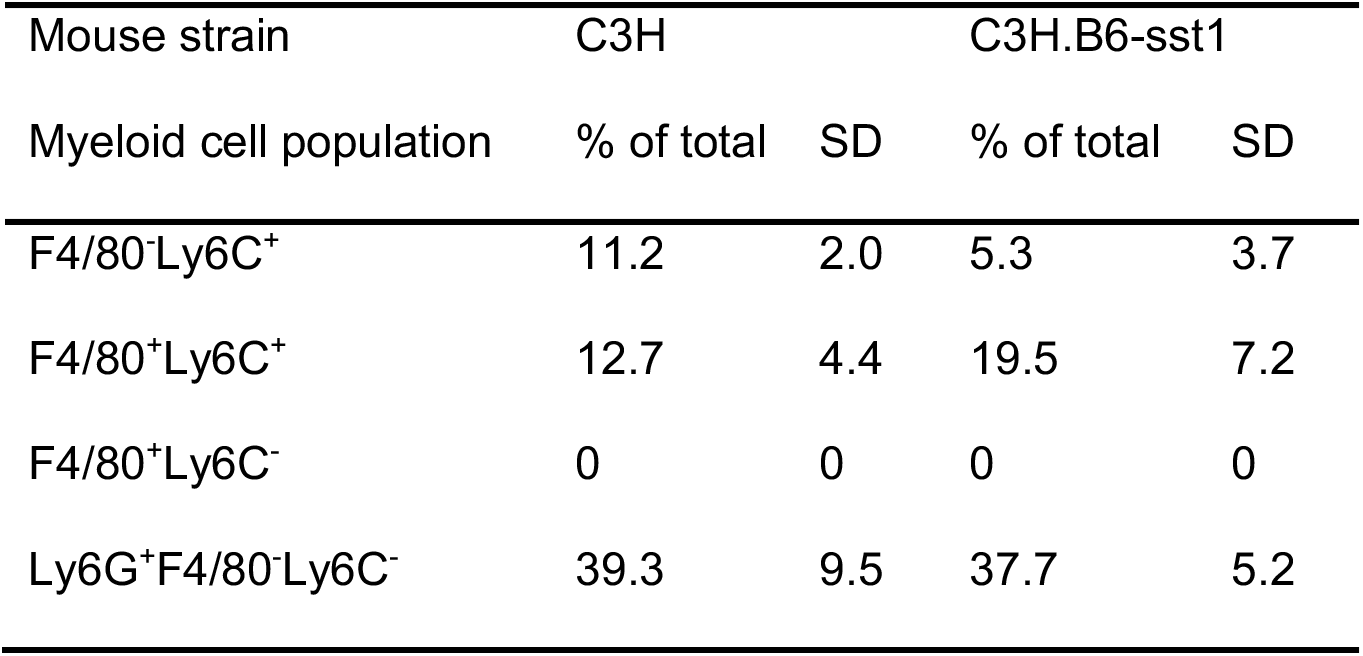
Flow cytometry of myeloid lung cells 8 days following aerosol challenge of C3H and C3H.B6-sst1 mice with 300 CFU of *F.t.* LVS

### Prolonged TNF stimulation induces bi-phasic progression of the integrated stress response and pro-apoptotic reprogramming in sst1S macrophages

Upon TNF stimulation, the Trib3 and Chac1 genes were among the most highly upregulated late response genes in the sst1S BMDMs. These genes are known targets of a transcription factor Chop10 (Ddit3), which is activated downstream of the ISR transcription factors ATF4 and ATF3 (**Suppl Fig. 1**). Collectively, this pathway is known as Integrated Stress Response (ISR)^22,23^. First, we compared the mRNA kinetics of genes representing transcriptional targets of the ISR (Atf3, Chop10, Chac1, Trb3, and Ddit4). The expression of the ISR genes spiked in the B6-sst1S cells at 16 h and continued to increase further between 16 – 24 h (**Fig. 2A**). Next, we monitored the expression of ISR markers ATF4 and ATF3 at the protein level by Western blot. Initially, we observed similar induction of ATF4 and ATF3 after 3 h of TNF stimulation in both the B6-sst1R and B6-sst1S BMDMs. However, in the B6 cells the levels of ATF4 and ATF3 proteins declined to basal levels by 15 and 24 h, respectively. Meanwhile, in the susceptible macrophages the ATF3 levels increased during the 16 – 24 h interval (**Fig. 2B**). Thus, the sst1S allele is uniquely associated with escalated transcription and translation of ISR genes after 12 hrs of TNF stimulation, timing which closely resembles the kinetics of the IFN-I pathway induction by TNF in sst1S macrophages.

**Figure 2.**
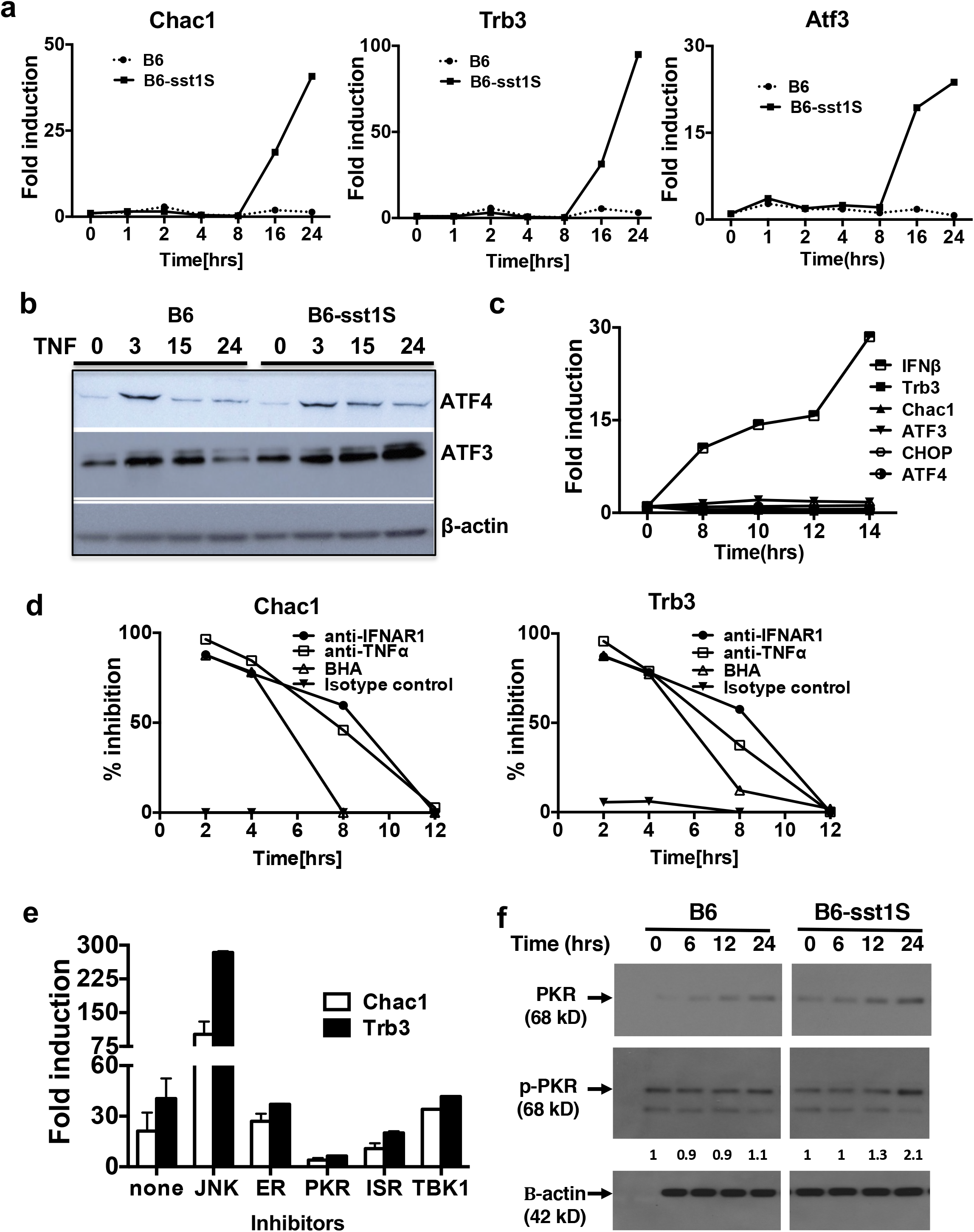
TNF treatment leads to bi-phasic upregulation of integrated stress response in B6-sst1S BMDM. **a)** Time course of the ISR gene (Chac1, T rb3 and Atf3) mRNA expression in B6-sst1 S and B6wt BMDMs within 24 h of treatment with TNF (10ng/mL). **b)** Timecourse of ISR protein expression in B6wt and B6-sst1S BMDMs stimulated with 10ng/mL of TNFa. ATF3 and ATF4 levels were determined using immunoblot in whole cell extracts (representative of two biological replicas). **c)** Time course of IFNβ and ISR gene expression in B6-sst1S BMDMs after stimulation with10ng/mL TNF for 8 -14 h. Real time PCR data is normalized to expression of 18S mRNA mRNA and presented relative to expression in untreated cells (set as 1). **d)** Time dependent effects of TNF and IFNAR blockade and ROS inhibition on Trb3 and Chac1 mRNA expression in B6-sst1S BMDM treated with 10ng/ml TNFa for 16hrs. BHA, a-IFNAR1, a-TNFa and isotype control antibodies were added after 2, 4, 8 and 12 h of TNF treatment. Trb3 and Chac1 mRNA expression was measured at 16 h of TNF stimulation and calculated as % inhibition with respect to cells treated with 10ng/mL TNFa alone. **e)** Effect of inhibitors on late phase ISR gene expression in TNF stimulated B6-sst1S BMDM. Inhibitors of JNK, ER stress (PBA), PKR, ISR and TBK1 were added after 12h of TNF stimulation (10 ng/mL), and Trb3 and Chac1 mRNA levels were measured at 16h. Trb3 and Chac1 mRNA expression was normalized to expression of 18S rRNA and presented relative to expression in untreated cells (set as 1). The qPCR results represent data from three independent experiments. **f)** Time course of PKR protein expression and phosphorylation (p-PKR) in B6-sst1S and B6wt BMDMs treated with TNF (10ng/mL) for indicated times. The Western blot is representative of two independent experiments. Numbers indicate fold induction of phospho-PKR, as compared to basal level (after normalization).

We followed the kinetics of the ISR- and IFN-inducible genes within a critical period between 8 and 14 hours at 2 h intervals. The IFNβ mRNA expression level in the B6-sst1S macrophages gradually increased, while the ISR markers remained at the same level throughout this period, suggesting a possible mechanistic hierarchy (**Fig. 2C**). Therefore, we tested whether blocking IFN-I signaling reduced the ISR induction. The IFN type I receptor (IFNAR1) blocking antibodies were added at different times after stimulation with TNF (10 ng/ml), and the ISR was assessed at 16 h of the TNF stimulation (**Fig. 2D**). Blocking the IFN-I signaling at 2 – 4 h after TNF stimulation profoundly suppressed the ISR escalation, as measured by Chac1 and Trb3 gene expression at 16 h of TNF stimulation. Blocking TNF signaling using neutralizing antibodies affected the ISR induction in a manner very similar to the IFNAR blockade – it was effective at the earlier stages (2-4 h) and only partially efficient at 8 h. Effects of both inhibitors on ISR completely disappeared by 12 h (**Fig. 2D**). Initially, ISR gene induction was prevented by ROS inhibitor BHA added at 2 – 4 h after TNF stimulation. However, its effect dramatically declined within 4 – 8 h of TNF stimulation. Thus, once the ISR pathway was set in motion, its transition from the latent to overt ISR in the sst1S macrophages after 12 h of TNF stimulation was driven in a TNF-, IFN- and ROS-independent manner.

The ISR is known to be induced as a result of the inhibition of cap-dependent translation caused by phosphorylation of eIF2α by several protein kinases activated in response to various stresses: viral infection (PKR), ER stress (PERK), starvation (GCN2), oxidative stress and hypoxia (HIPK)^24^. To reveal the driving force behind the late ISR transition in the sst1S macrophages, we measured the post-TNF induction of Trb3 and Chac1 mRNAs in B6-sst1S BMDM in the presence of small molecules which inhibit either the ISR (ISRIB, an eIF2α phosphorylation inhibitor)^25^, ER stress (PBA)^26^, PKR (C16)^27^, TBK1 (BX-795), and stress kinases p38 (SB203580) and JNK (SP600125)(**Fig. 2E**). Added at 12 h post TNF stimulation, the ISR and PKR inhibitors (ISRIB and C16, respectively) profoundly inhibited the upregulation of both sentinel mRNAs, while the ER stress inhibitor PBA and the TBK1 kinase inhibitor BX-795 had no effect. This suggests that PKR activity was responsible for transition from latent to overt ISR specifically in the B6-sst1S macrophages. Indeed, PKR levels increased by the 12 h of TNF stimulation and were subsequently maintained at higher level in the B6-sst1S cells. The level of active, phosphorylated, PKR also increased 12 – 24 h of TNF stimulation exclusively in B6-sst1S macrophages (**Fig. 2F**). PKR is a classical interferon-inducible protein, whose kinase activity is induced by double-stranded RNA (dsRNA). Traditionally, the PKR pathway was associated with antiviral immunity, but more recently it has been demonstrated that, in addition to viral dsRNAs, PKR can interact with and be activated by misfolded and dimerized endogenous RNA molecules^28,29^. Using dsRNA-specific antibodies J2^30^, we detected dsRNA speckles in the cytoplasm of TNF-stimulated BMDM of both B6 and B6-sst1S backgrounds (**Suppl. Fig.4A**). Thus, the presence of the endogenous PKR ligands may provide an explanation of how IFN-induced PKR is activated by TNF in non-infected macrophages. The PKR activation by the endogeneous ligands has been linked to metabolic dysregulation via activation of stress kinase JNK^31^. We observed that JNK inhibitor increased the expression of the ISR markers (**Fig. 2E**) suggesting a role for the PKR – JNK-mediated feedback circuit proposed by Nakamura et al. in our model (see below).

These data demonstrate that in sst1S macrophages TNF initiated a cascade of stress responses in biphasic manner. The early initiation phase (2 – 4 hrs) required ROS, leading to ER stress and ER stress-mediated ISR similarly in both the wt and sst1S mutant macrophages. In support of this notion, the levels of classical ER stress marker BiP were induced to similar levels in TNF-stimulated B6wt and B6-sst1S macrophages. Also, XBP-1 splicing, known to be induced by IRE1 kinase activated specifically by the ER stress follows similar kinetics in both strains reaching peaks at 4 – 8 h (**Suppl. Fig.4B**), corresponding to early phase ATF4 protein upregulation (**Fig. 2B, above**). While self-limited in the B6wt cells, the ISR escalated in the B6-sst1S macrophages during the 12 – 24 h period of TNF stimulation via distinct, IFN-I-dependent, mechanism involving PKR activation.

The major adaptive role of ISR is a global reduction of cap-dependent protein translation^22^. However, translation of many proteins involved in stress responses proceeds via cap-independent mechanisms, and the proportion of those proteins in the cellular proteome increases during prolonged stress. Thus, we postulated that an unrelenting stress in B6-sst1S macrophages would result in global proteome remodeling, and discordance between transcriptome and proteome. Indeed, we detected upregulation of Trb3 protein at the early stage of TNF stimulation, but not during the second phase of ISR escalation in B6-sst1S macrophages, despite high levels of its mRNA induction (**Suppl. Fig. 5A**). Therefore, we compared global quantitative protein abundance profiles of the B6wt and B6-sst1S mutant macrophages after stimulation with TNF, using stable isotope labeling of the digested macrophage proteomes with tandem mass tags (TMT) followed by deep 2-D LC-MS/MS-based proteomic analysis.

The proteome profiles of TNF-stimulated B6-sst1S and B6wt macrophages were clearly distinct (**Suppl. Table 1**). A number of proteins were up-regulated in the susceptible macrophages demonstrating the absence of a total translational arrest in the mutant cells. In agreement with Hspa1a mRNA induction (**Fig. 1E**), higher levels of Hspa1a protein were detected by the proteomic and Western blot analyses (**Suppl. Fig. 5B**). We also observed that the TNF-stimulated mutant sst1S cells expressed higher levels of proapototic proteins DAXX and Bim, cold shock-inducible RNA binding protein Rbm3 and dsRNA-binding protein Stauphen1. We confirmed the upregulation of DAXX and Bim by Western blot (**Suppl. Fig. 5C-D**). In contrast, the proteome of B6wt macrophages stimulated with TNF was enriched in proteins involved in antioxidant defenses and protein homeostasis in ER and cytoplasm, such as: 1) NADH-cytochrome b5 reductase 4 (CYB5R4) which protects cells from excess buildup of ROS and oxidant stress^32^; 2) Stromal cell-derived factor 2 (SDF2), involved in ER protein quality control, unfolded protein response and cell survival under ER stress^33^; 3) The signal sequence receptor 2 (SSR2), a subunit of the ER TRAP complex involved in protein translocation across the ER membrane^34^; 4) Stress-associated endoplasmic reticulum protein 1 (SERP1) which interacts with target proteins during their translocation into the lumen of the endoplasmic reticulum and protects unfolded target proteins against degradation during ER stress^35^. Only the B6wt cells expressed the SP110/IPR1 protein encoded within the *sst1* locus. Ipr1 has been identified and validated as a strong candidate gene in our previous work using positional cloning^17^. The IPR1 protein was present in proteome of both non-stimulated and, at higher levels, of TNF-stimulated B6 macrophages. We extended these observations by finding that the IPR1 protein was induced in B6wt macrophages between 8 and 12 h after initial TNF stimulation, corresponding to a period of late stress escalation in the IPR1 – negative B6-sst1S cells (**Suppl. Fig. 5G**). Inhibition JNK, p38 or IFNAR blockade prevented the IPR1 protein up-regulation by TNF demonstrating that it is an interferon- and stress-inducible protein (**Suppl. Fig. 5H**).

Compared to the B6wt, TNF-stimulated B6-sst1S macrophages also expressed lower levels of cyclin-dependent kinase inhibitor p21 and 7-dehydrocholesterol reductase (Dhcr7), a key enzyme in cholesterol biosynthesis. Using Western blot we determined that the difference was due to the down-regulation of p21 and Dhcr7 in the B6-sst1S macrophages 12 – 16 h of TNF stimulation (**Suppl. Fig. 5E-F**). The Dhcr7 downregulation might be explained by the inhibitory effect of 25-hydoxycholesterol on cholesterol biosynthesis via SREBP2 inhibition^36^. This oxidized cholesterol derivative is produced by the IFN-I-inducible enzyme Ch25h (cholesterol 25-hydroxylase)^37^, which is highly upregulated in the B6-sst1S macrophages by TNF (**Fig. 1E**).

Taken together the above data demonstrate that compromised stress resilience of the sst1S macrophages after TNF stimulation is mechanistically linked to hyperactivity of IFN-I pathway, PKR-sustained ISR escalation, pro-apoptotic proteome remodeling and, possibly, broader down regulation of metabolic pathways, such as cholesterol biosynthesis, in IFN-I-dependent manner. Therefore, we wanted to test whether inhibitors of IFN-I and ISR pathways can correct the susceptible macrophage phenotype and, thus, decrease susceptibility to virulent Mtb in vivo.

### Small molecule inhibition of the ISR reduces susceptibility to Mtb and granuloma necrosis in vivo

To test the in vivo role of TNF-mediated IFN-I and ISR hyperstimulation, we tested small molecule inhibitors of these pathways in B6-sst1S mice in a model of chronic tuberculosis. First we evaluated ISRIB, an inhibitor of eIF2α phosphorylation which has been shown to reduce the ISR^38^. Four weeks after aerosol infection with Mtb H37Rv, groups of mice were treated daily (5/7 days) for 8 weeks with ISRIB at 0.25 mg/kg, 1.0 mg/kg or with vehicle control (**Suppl. Fig. 6A**). Low-dose ISRIB, and to a lesser extent high-dose ISRIB, statistically significantly inhibited bacterial proliferation (**Fig. 3A**) and the degree of focal pneumonia with granuloma formation (**Fig. 3B, Suppl. Fig. 6B**). Two-dimensional evaluation of lung pathology with digitally scanned sections collected at both 4 wk and 8 wk following the initiation of therapy revealed that both doses of ISRIB significantly reduced lung inflammation and necrosis (**Fig. 3C, Suppl. Fig 6C**). To achieve a more global assessment of the impact of ISRIB therapy, we used three-dimensional scanning with CT and [^18^F]FMISO PET in live mice at 8 weeks of therapy. Quantitative, unbiased evaluation of non-vessel, non-airway, lung voxels associated with lung tissue consolidation revealed a trend towards reduced lung consolidation in the ISRIB-treated mice compared with untreated. (**Fig. 3D**).

**Figure 3.**
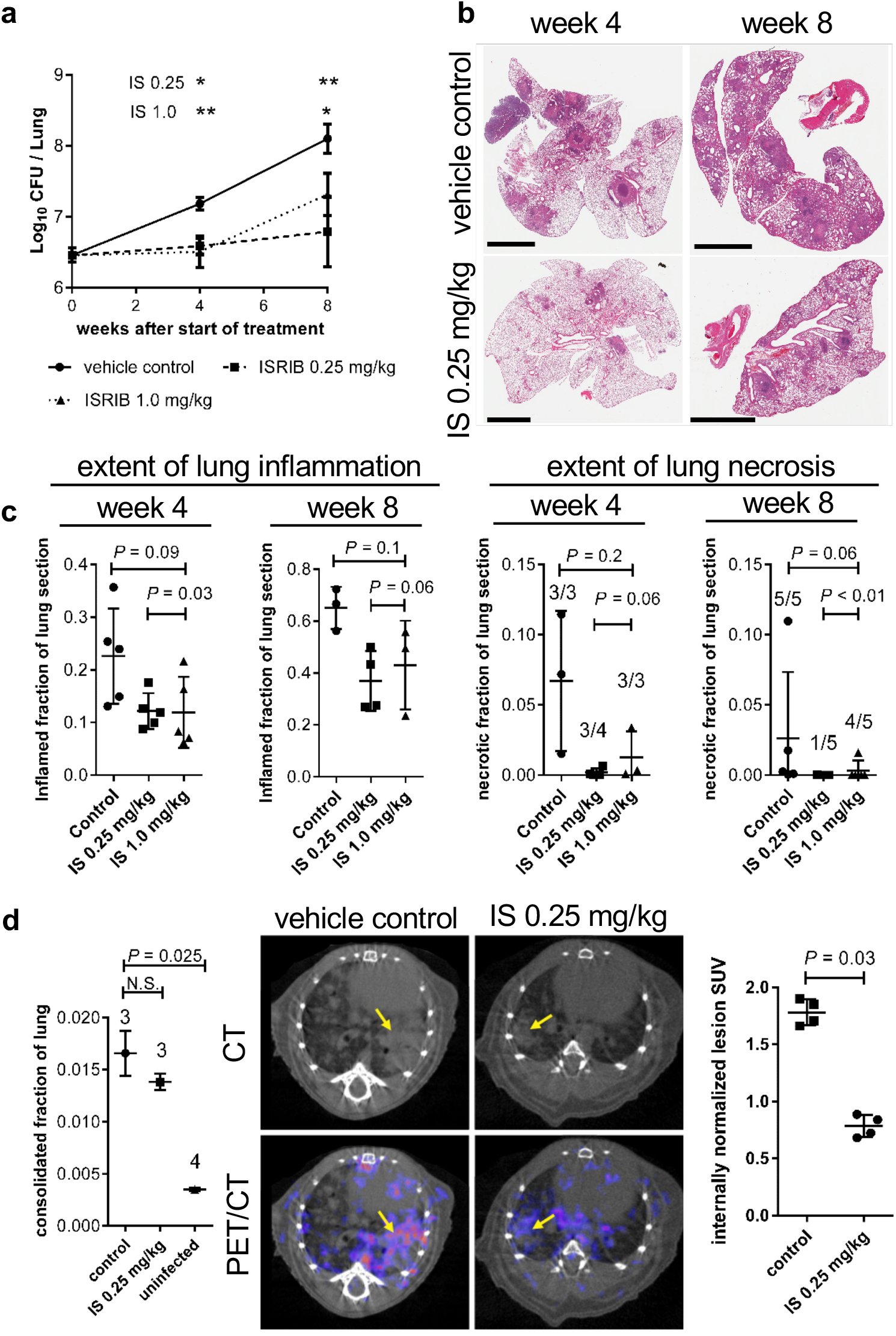
Effect of ISR inhibitor ISRIB on tuberculosis progression in B6-sst1S mice. For all figure panes, IS=ISRIB. **a)** Lung bacillary loads at weeks 4 and 8 after treatment start. Data plotted as means±SEM. Asterisks indicate significant differences between treatment groups compared to vehicle control group calculated by two-way ANOVA. * indicates significance at 90% CI. ** indicates significance at 95% CI. **b)** Representative H&E stained lung tissue slices from mice in the vehicle control group and the ISRIB treated group with 0.25 mg/kg by body weight. Scale bars = 3 mm. **c)** Extent of inflammation and necrosis in lungs of mice treated with vehicle control or ISRIB. Each data-point indicates a survey of one mouse. All P-values calculated based on a non-parametric two-tailed Mann-Whitney test. Fractions above group columns indicate the numbers of mice where necrosis was identified / the number of mice in analysis group. **d)** Fraction of lung volume with disease density as a quantitative measure of severity of disease in mouse lungs at week 8. Data summary elements represent mean fraction of voxels±SEM. *P*-values calculated based on Welch’s t-test. N.S.= not significant. Numbers above group columns indicate number of mice surveyed. **e)** [^18^F]FMISO PET/CT imaging of mouse lungs at week 8. Quantification is given as dose and decay corrected standardized uptake value (SUV) of lesions normalized to PET signal from PET-blinded CT-scan selection of non-disease lung space volumes. P-values calculated by non-parametric two-tailed Mann-Whitney test. For charts c and e, chart data summary elements indicate means±SD.

We also assessed the extent of hypoxia within the cells of inflammatory foci since intracellular hypoxia is a known factor linking the ISR to the onset of necrosis^39^. We used lesion-specific [^18^F]FMISO PET signals from our CT/PET scans since [^18^F]FMISO is known to accumulate in the cytoplasm of hypoxic cells^40,41^. Mice treated with the ISR inhibitor showed a >2-fold reduction in mean internally normalized [^18^F]FMISO lung lesion SUV signal compared with controls supporting the conclusion that lesions from ISRIB treated mice contained intracellular conditions that were less disposed to the onset of necrosis. (**Fig. 3E**). We also tested ISRIB for direct antibacterial activity against Mtb broth cultures and found no significant activity with an MIC >256 μg/ml. On testing the ability of ISRIB to inhibit the proliferation of Mtb in B6-sst1S BMDM, we found no antibacterial effect (**Suppl. Fig. 6D**) confirming that that ISRIB does not directly activate macrophage killing of Mtb, but rather its actions in blunting macrophage integrated stress response pathway in vivo promote balanced inflammatory and antimicrobial macrophage responses and prevent lung tissue damage.

Next, we tested effect of ISRIB in a mouse model of more severe TB using C3HeB/FeJ mice that carry the sst1S allele on a more susceptible to Mtb genetic background^42^(**Suppl. Fig. 7A**). In this model, low-dose ISRIB again revealed potent effect significantly reducing lung Mtb burdens (**Suppl. Fig. 7B**). In parallel, we assessed the mTor pathway inhibition in this same mouse TB model, since this pathway has been reported to downregulate the mitochondrial integrated stress response^43^ and to induce autophagy^44^. However, treatment with the mTor inhibitor sirolimus was ineffective (**Suppl. Fig. 7D**). Because the sst1S-mediated ISR is IFN-I driven, we tested a potent IFI-I pathway inhibitor amlexanox, which prevents phosphorylation of TBK1 and IKKe. Unexpectedly, amlexanox at both 25 mg/kg and 100 mg/kg was inactive or even deleterious in terms of controlling lung Mtb loads and lung pathology (**Suppl. Fig 7C and 7E**, respectively). This result suggests that sst1S-mediated susceptibility is independent of the TBK1-medited pathway. It is also consistent with our in vitro experiments demonstrating that TNF-induced ISR escalation in sst1S macrophages is IFN-I driven, but TBK1-independent (**Fig. 2E**).

### JNK and Myc hyperactivity underlie the IFNβ super-induction in TBK1-independent manner

A previous report demonstrated that in wild type B6 macrophages, TNF stimulated low levels of IFNβ via NF-kB-mediated induction of IRF1, followed by auto-amplification by secreted IFNβ via the IFN-I receptor and IRF7^21^. In our model, the IRF1 protein was similarly upregulated by TNF in both B6wt and B6-sst1S mutant macrophages (**Fig. 4A**). To determine which of the IRF transcription factors might play a dominant role in the IFN-I pathway hyper-activation observed in the *sst1* susceptible macrophages, we performed knockdowns of IRF1, IRF3 and IRF7 prior to stimulation of BMDMs with TNF using siRNAs (**Fig. 4A and Suppl. Fig.8A and 8B**). The IRF1 knockdown had the most pronounced effect, while the IRF3 and IRF7 knockdowns had similar but weaker effects on IFNβ mRNA expression after 16 h of TNF stimulation (**Fig. 4B**). Importantly, knockdowns of any of those IRF1s reduced the IFNβ expression proportionally in both wt and mutant macrophages and did not eliminate the strain difference. Also, IFNAR1 blocking antibodies were ineffective in preventing the late phase IFNβ upregulation in the B6-sst1S cells when added after 8 hr of TNF stimulation, while TNF blockade remained efficient (**Fig. 4C**) indicating that the late IFNβ upregulation in the sst1S background was not due to autoamplification by secreted IFNβ.

**Figure 4.**
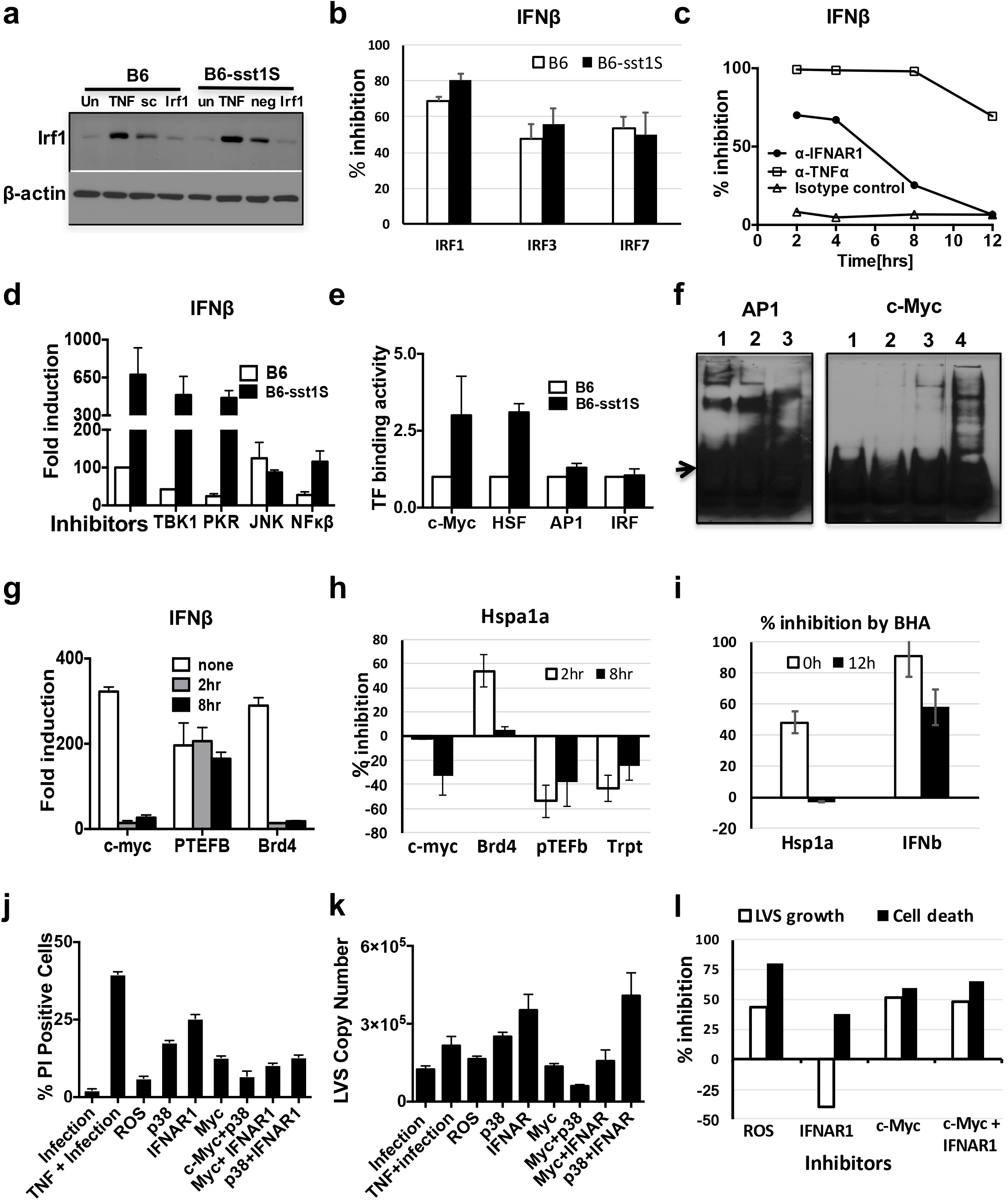
Transcriptional control of IFNβ super-induction in B6-sst1S macrophages by TNF. **a)** Effects of TNF stimulation and siRNA knockdown on IRF1 protein expression in B6 and B6-sst1S BMDMs stimulated with 10 ng/ml of TNF for 24 h. Cell were treated with siRNA 24 h prior to stimulation with TNF. Immunoblot using IRF1-specific polyclonal antibodies represents data from two independent experiments. **b)** Inhibition of IFNβmRNA expression in TNF-stimulated B6 and B6-sst1S BMDM after knockdown of IRF1, IRF3 and IRF7 using siRNA. Percent inhibition was calculated as compared to scrambled siRNA control. The data are representative of three independent experiments. **c)** Effect of TNF and IFN-I receptor blockade on IFNβ mRNA expression in B6-sst1S BMDM treated with 10ng/ml TNFa for 16 h. a-IFNAR1, a-TNFa and isotype control antibodies (10 ug/ml) were added at 2,4, 8 and 12 h of TNF treatment. Percent inhibition of IFNβ mRNA expression was calculated as compared to cells treated with 10ng/mL TNFa and isotype control antibodies at 2 h. The data are representative of two independent experiments. **d)** Effect of small molecule inhibitors on IFNβmRNA expression in B6-sst1S BMDM treated with 10ng/ml TNFa for 16 h. Inhibitors of TBK1, PKR, JNK and NF-ĸB were added after 12 h of TNF stimulation (for four hours). IFNβ mRNA expression was measured using qRT-PCR and normalized to expression of 18S rRNA. The fold induction of gene expression is calculated relative to the mRNA expression in untreated cells (set as 1). The data are representative of two independent experiments. **e)** Transcription factor (TF) binding activities were compared using Transcription Factor Profiling Array (Signosis). Nuclear extracts (NE) were isolated from B6 and B6-sst1S BMDMs stimulated with TNF (10 ng/ml) for 12 h. Results represent data from two independent experiments. **f)** Validation of TF arrays results using EMSA with c-Myc and AP-1 probes. Nuclear extracts (NE) were isolated at 12 h of TNF stimulation, as above. Left panel – AP-1: 1-B6wt NE, 2 – B6-sst1S NE, 3-cold probe competition. Right panel – c-Myc: 1-free probe, 2 – B6wt NE, 3 – B6-sst1S NE, 4-cold probe competition. The arrow denotes free probes. Results represent data from two independent experiments. **g)** Effect of inhibitors on IFNβ mRNA expression in TNF stimulated B6-sst1S BMDM. Inhibitors of c-Myc (10058-F4), pTEFb (Flavopiridol) and Brd4 (JQ1) were added after 2h and 8h of TNF stimulation. IFNβ mRNA expression was measured at 18 h of stimulation with TNF (10ng/mL). **h)** Effect of transcription inhibitors on Hspa1a mRNA expression. Hspa1a mRNA expression was measured after 18 h of stimulation with 10 ng/ml of TNF. Inhibitors of c-Myc (10058-F4), pTEFb (Flavopiridol), Brd4 (JQ1) and RNA pol II inhibitor Triptolide (Trpt) were added after 2h and 8h of TNF stimulation. **i)** Effects of an oxidative stress inhibitor BHA on TNF-induced Hspa1a and IFNβ mRNA levels in B6-sst1S BMDMs. BHA was added before or 12 h after TNF stimulation. The mRNA levels were determined using qRT-PCR at 18 h of TNF stimulation. % inhibition was calculated as compared to TNF stimulated cells not treated with BHA. **j-k)** Effects of inhibitors and their pairwise combinations on macrophage cell death (j) and bacterial loads **(k)** in TNF-stimulated B6-sst1S BMDM infected with F.t.LVS (MOI=1). Cells were primed with TNF (10 ng/ml) for 16 h and subsequently infected with *F.t.* LVS at MOI=1 for 24 hrs. The inhibitors were added after 4hrs of TNF treatment. Cell death (j) was measured as % of PI positive cells using automated microscopy. The bacterial enumeration **(k)** was done using real time qPCR and was normalized to total cell number determined using automated microscopy (Celigo). The bacterial load and cell death data are representative of three independent experiments. **l)** Comparative effects of ROS, IFN-I and c-Myc inhibition on the *F.t.* LVS-infected macrophage death and the bacterial control summarized from panels j and k.

To identify pathway(s) specifically responsible for the late stage IFNβ super-induction in the B6-sst1S macrophages, we used small molecule kinase inhibitors. We added these agents after 12 h of TNF stimulation, and measured the IFNβ mRNA levels four hours later (**Fig. 4D**). Unsurprisingly, NF-kB inhibitor (BAY11-7082) proportionally reduced the IFNβ mRNA levels in both wt and mutant macrophages. Strikingly, inhibiting JNK completely eliminated the *sst1-* dependent difference: JNK inhibitor SP600125 reduced the IFNβ mRNA expression in the B6-sst1S macrophages to the level of B6, but did not affect the IFNβ expression level in B6 macrophages. In contrast, inhibition of another IFN-inducing kinase TBK1, which is involved in signaling by nucleic acid recognition modules and IRF3 activation, did not affect the IFNβ induction in either wt or the sst1S cells (**Fig. 4D**). These findings reveal that the *sst1* locus neither acted by effects on the canonical TNF – IRF1 – IFNβ axis and the IFNβ – IFNAR1 – IRF7 – IFNβ auto-amplification loop, nor did it control the TBK1 – IRF3 pathway. Rather, the late phase superinduction of IFNβ, exclusively observed in *sst1S* macrophages, results from stress-activated kinase JNK cooperation with the canonical TNF – NF-kB – IRF1 pathway.

To gain deeper insight into *sst1*-mediated transcriptional regulation at this critical transition period, we compared transcription factor (TF) activities in B6wt and B6-sst1S macrophages following 12h of TNF stimulation using a TF activation array (Signosis). The activities of NF-kB, AP1, STAT1, GAS/ISRE, IRF, NFAT, NFE2, CREB, YY1 and SP1 were upregulated by TNF to a similar degree in the wt and mutant macrophages, while the HSF1 and MYC consensus sequence binding was significantly upregulated only in the mutant cells (**Fig. 4E**). The higher TF activity of Myc in TNF-stimulated B6-sst1S macrophages was confirmed using EMSA (**Fig. 4F**). Next, we tested whether increased c-Myc activity in susceptible macrophages amplified IFNβ transcription. Indeed, knockdown of c-Myc using siRNA significantly reduced IFNβ mRNA expression (**Suppl. Fig 9A**). The small molecule inhibitor of Myc-Max dimerization 10058-F4, which suppresses E box – specific transcriptional activation by this heterodimer, i.e. targets promoter-specific TF activity of c-Myc, potently suppressed TNF-induced IFNβ mRNA expression (**Fig. 4G**). Myc is also known to promote activity of transcriptionally active genes independently of its binding to specific promoters^45,46^. Therefore, we tested whether the inhibitor of positive transcription elongation factor (p-TEFb) flavopiridol or the RNA Pol II inhibitor of transcriptional elongation triptolide could also inhibit the IFNβ super-induction and found that both of them were inactive. In contrast, the Brd4 inhibitor JQ1, which has been previously shown to directly suppress Myc-dependent transcription^47^, was as active as 10058-F4 (**Fig. 4G**). These data demonstrate that increased activity of c-Myc transcriptional network significantly contributes to the IFN-I pathway hyperactivation.

Using TF Array, we also observed increased HSF1 activity, which is consistent with escalating Hspa1a protein and mRNA expression in TNF-stimulated B6-sst1S macrophages (**Suppl. Fig.5B and 9B**, respectively). The HSF1 TF activation and heat shock protein induction by TNF are indicative of proteotoxic stress (PS). In the wild type cells the heat shock stress response was moderate and did not escalate past 12 h, while in the susceptible cells it significantly increased between 8 and 16 h of TNF stimulation. The HSF1 inhibitor KRIB11, which blocks HSF1 activity, induced death of TNF-stimulated macrophages irrespective of their *sst1* genotype (**Suppl. Fig.9C**), demonstrating that, initially, PS was experienced by TNF stimulated macrophages of both backgrounds and the HSF1-mediated stress response was an important survival pathway. Remarkably, another HSF1 inhibitor RHT^48^ that, in addition to blocking HSF1 also reduces protein translation, suppressed Hspa1a mRNA expression without killing macrophages and eliminated the difference between the wt and sst1S macrophages in IFNβ mRNA expression, when added 2h after TNF stimulation(**Suppl. Fig.9D**). In contrast to the IFNβ regulation, inhibition of c-Myc did not prevent late PS escalation in the sst1S cells (**Fig. 4H and Suppl. Fig.9E**). We concluded that greater PS/HSF1 pathway activation in sst1S macrophages by TNF occurred either in parallel or upstream of the IFNβ super-induction. Indeed, PS is known to induce JNK activation that is a driving force behind the IFNβ super-induction seen in our model.

Reactive oxygen species (ROS) are known to cause protein misfolding and aggregation in cytoplasm and ER. We found that pre-treatment of B6-sst1S macrophages with BHA to boost their anti-oxidant defenses prior to TNF stimulation inhibited super-induction of Hspa1a and IFNβ (**Fig. 4I**) and prevented subsequent escalation of the ISR (**Suppl. Fig. 9F**). The levels of ROS produced by TNF-stimulated B6wt and B6-sst1S macrophages were similar (**Suppl.Fig.9G**) suggesting that those cells differ in adaptation to ROS-mediated stress rather than overall ROS levels. Taken together, these data demonstrate that an aberrant response of the *sst1S* macrophages to TNF is initiated by ROS and characterized by intensifying proteotoxic stress while c-Myc activity remains elevated. Both pathways contribute to further stress escalation via superinduction of the type I IFN pathway and transition to overt, unchecked upregulation of the ISR.

To assess the impact of individual stress pathways on infected macrophage survival and bacterial control, we pre-treated B6-sst1S BMDMs with TNF in the presence of ROS, p38, c-Myc inhibitors and IFNAR blocking antibodies, as well as their pairwise combinations. Then the cells were infected with *F.t.* LVS at MOI 1:1 and 3:1. Inhibitors of ROS (BHA), p38, c-Myc, as well as IFNAR blocking antibodies, significantly reduced cell death (**Fig. 4J**). Next, we studied effects of inhibitors on the bacterial control. The c-Myc inhibitor significantly reduced, while the IFNAR1 receptor blockade increased, the bacterial loads in TNF treated B6-sst1S macrophages, while ROS and p38 inhibitors were neutral in this respect (**Fig. 4K**). Overall, these data demonstrate that the same mechanisms that underlie TNF-induced stress also compromise survival and bacterial control of the infected macrophages. Interestingly, inhibition of Myc transcription factor activity uniquely improved both macrophage survival and bacterial control, while inhibition of IFNAR1 only improved macrophage survival, but reduced the bacterial control (**Fig. 4L**). Remarkably, in pairwise combinations, the c-Myc inhibitor could overcome detrimental effect of the IFNAR blockade on the bacterial replication in agreement with our previous data demonstrating that Myc hyperactivity mechanistically is upstream of IFNβ super-induction and ISR activation in B6-sst1S macrophages.

## DISCUSSION

Taken together, the complex cascade delineated in this study indicates that the *sst1-* mediated susceptibility to intracellular bacterial pathogens is mechanistically linked to an aberrant macrophage response to TNF and sustained escalating stress responses. IFN type I plays an important intermediary role in this cascade, with its super-induction resulting from a synergistic effect of the canonical NF-kB-mediated TNF activation pathway and a sst1-specific response mediated by the stress kinase JNK. The origins of stress, however, are upstream of the IFN-I pathway and reflect hyperactivity of Myc in the setting of TNF-induced oxidative stress. Inhibition of either transcriptional activity of Myc or ROS generation prevented the late phase IFN-I superinduction and the escalation of the ISR in TNF-stimulated sst1S macrophages. Thus, we suggest that deciphering stress-mediated regulation of Myc activity in macrophages is key to understanding the sst1-mediated phenotype in vitro and in vivo.

The *sst1* locus encodes interferon-inducible nuclear protein Sp110, whose expression is completely abolished in TNF- or IFNγ-activated macrophages carrying the *sst1* susceptible allele^17^. In the wild type macrophages, this protein is induced at the late stage of TNF response between 8 – 12 h. This time interval is critical for the expression of the susceptible phenotype, because it precedes the escalation of the proteotoxic stress and super-induction of IFNβ. During this period, Sp110 accumulates in the nucleus and associates with chromatin (Bhattacharya and Siedel, unpublished). We hypothesize that this protein may play direct or indirect roles in macrophage stress resilience via controlling chromatin organization and function.

Recent studies revealed a central role of another protein of the Sp100 family also encoded within the sst1 locus in macrophage activation: an interferon-inducible chromatin-binding bromodomain protein Sp140^49^. They have shown that in activated macrophages the SP140 protein plays an important role in preserving macrophage-specific transcriptional program by binding to promoters of “lineage-inappropriate” repressed genes and maintaining their repressed status. Among those was the developmental HOXA9 gene whose activity preserves hematopoietic stem cell self-renewal and suppresses macrophage differentiation. The Sp140 knockdown led to derepression of HOXA9 gene and aberrant macrophage activation including an upregulation of Myc and E2F regulated gene sets. This suggest that defect in bromodomain-mediated transcriptional repression may contribute to the *sst1* susceptible phenotype. Indeed, we observed that a small molecule bromodomain inhibitor JQ1^47^ repressed the IFNβ super-induction by TNF in sst1S macrophages in vitro.

In vivo, Sp140 downregulation resulted in exacerbated inflammatory colitis. In humans, Sp140 polymorphisms are associated with Crohn’s disease and multiple sclerosis in GWAS studies. The Sp110 polymorphisms have been associated with severity of canine degenerative myelopathy, a neurodegenerative disease with similarities to human ALS^50^. We hypothesize that in activated macrophages the Sp100 family members play a central role in cross talk of stress-, Myc- and interferon-mediated pathways to maintain macrophage differentiation and activation programs and boost macrophage stress resilience. This paradigm would explain broad roles of those proteins in limiting inflammatory tissue damage initiated by infectious and non-infectious triggers. Thus, the molecular mechanisms mediated by genes encoded within the *sst1* locus, as well as their human homologs, in activated macrophages remain high priority to be elucidated.

While Myc activity is important for growth of myeloid precursors^51^ and alternative macrophage activation in tumor microenvironment^52^, down regulation of Myc activity in monocytes/macrophages appears to be an important adaptive mechanism within infection-induced inflammatory lesions. The inflammatory monocytes are produced from myeloid precursors in bone marrow and recruited from circulation to sites of inflammation where they undergo final cell divisions and terminal differentiation. Prior to pathogen encounter, their phenotype is further sculpted by multiple factors within the inflammatory milieu, which include not only classic inflammatory mediators, but also growth factors, hypoxia, starvation, acidosis, etc.

Sensing and integrating responses to gradients of those factors is necessary to optimally balance cell metabolism, growth and macrophage effector functions within specific environments^8^. Indeed, in non-transformed cells, sensing stress can stop cell cycle progression and trigger terminal differentiation^53^. For example, stress kinases can stimulate p53 to block transcription of c-Myc and its targets^54^. Considering macrophage maturation within inflammatory lesions, this response may dramatically reduce energy expenditure and prepare macrophages for subsequent stress escalation and pathogen encounter. Indeed, we determined that Myc transcriptional activity was down regulated 12 hours after TNF stimulation, but only in the wild type macrophages. In contrast, the inability to repress Myc in TNF-stimulated susceptible macrophages is associated with escalation of proteotoxic stress (PS), possibly due to misfolding of newly synthesized proteins caused by ROS. Thus, persistent transcriptional activity of Myc appears to be a major driver of the maladaptive response of the sst1-susceptible macrophages, leading to un-resolving stress, accelerated death of infected macrophages and, eventually, to the development of overt necrotic pathology.

Our studies revealed the dynamics of downstream cascade associated with the susceptible phenotype. At an early stage, TNF stimulation causes proteotoxic stress (PS) and integrated stress response (ISR) in both wild type and the sst1-susceptible mutant macrophages. At this stage the ISR is driven by the ER stress and unfolded protein response (UPR), as previously described^55^. The unusual, second wave of ISR activation is exclusive for the sst1-susceptible phenotype. It is initiated by TNF in IFN-I-dependent manner via PKR activation, similar to a cascade recently described in a model of *Listeria monocytogenes* infection^56^. However, in the sst1-susceptible macrophages, the IFNβ super-induction was triggered by TNF alone. We excluded a significant contribution of the STING – TBK1 – IRF3 pathway to the observed IFNβ super-induction and, thus, demonstrated that recognition of endogenous or exogenous nucleic acids was not required. Instead, the upregulation of the IFN-I pathway could be explained solely by a cooperative effect of prolonged activation of NF-ĸB and JNK pathways. Both pathways are known to converge on IFNβ enhancer and recruit coactivators and chromatin-remodeling proteins to form an enhanceosome^57,58^. JNK is activated in response to oxidative, proteotoxic, metabolic and other challenges and is an important part of the cellular defense strategy against stress^59,60^. Whereas transient JNK activation is adaptive, prolonged JNK activation is known to contribute to pro-apoptotic transition, which, as we show, in TNF-stimulated sst1-susceptible macrophages occurs via type I IFN pathway upregulation and PKR activation.

Numerous studies reveal a role of the type I interferon (IFN-I) pathway in the immunopathology caused by intracellular bacteria, including Mtb^61–64^, chronic viral infections^65–67^ and autoimmunity (reviewed in^68–70^). Our studies reveal a mechanism of stress-mediated IFN-I pathway upregulation that makes macrophages less resilient to subsequent infection with intracellular bacteria and is associated with immunopathology in vivo. We propose that stress-induced IFN-I pathway escalation mechanistically represents an intermediate step between a low grade constitutive IFN-I pathway activity that plays homeostatic role^71^ and a full blown activation downstream of nucleic acid recognition receptors mediated by IRF3. Normally, it plays an adaptive role. For example, upregulation of JNK by proteotoxic stress induced by TNF may lead to transient activation of the IFNβ – PKR – ISR axis to limit global protein biosynthesis by inhibiting cap-dependent translation. However, un-resolving proteotoxic stress and a persistently elevated ISR observed in susceptible macrophages leads to further escalation of JNK activity, corresponding increase in IFNβ production and upregulation of interferon-stimulated genes PKR, Rsad2 and Ch25h, whose products inhibit protein biosynthesis, mitochondrial function and lipogenesis, respectively^56,72,73^. The 25-hydrocholesterol produced by the Ch25h enzymatic activity can further increase the ISR^74^ and amplify inflammatory cytokine production^75^. Moreover, by limiting cholesterol biosynthesis it can sustain elevated IFN-I signaling^76^. Previously, PKR has been shown to stimulate JNK activity in macrophages^31^. Possibly, this occurs via translational arrest, since other protein synthesis inhibitors also activate stress kinases JNK and p38 via a mechanism known as the “ribotoxic stress response”^77,78^. Thus, it appears that at certain levels JNK, IFNβ and PKR may form a maladaptive feed-forward stress response circuit, locking TNF-stimulated sst1-susceptible macrophages in a state of unrelenting stress and, eventually, leading to an IFN-I-dominated hyper-inflammatory response, suppression of essential metabolic pathways and, eventually, macrophage damage.

Obviously, this susceptibility-associated mechanism represents an attractive therapeutic target. Indeed, this paper reveals that a small molecule inhibitor of the ISR, ISRIB, is effective in preventing necrosis in Mtb-infected mouse lungs and concomitantly restricts Mtb proliferation. ISRIB has demonstrated promising results in prevention of neuronal degeneration, memory enhancement, and reducing tumor growth in certain cancers^79,80^. Further studies in combination with traditional anti-TB drugs may define whether future ISR inhibitors which are being developed for human use may offer therapeutic benefit as a host-directed therapy for TB and related intracellular infections. On a cautionary note, however, elements of this pathway may represent imperfect, but necessary, backup strategy of stress adaptation, and their inhibition may be detrimental. Thus, we observed that inhibition of HSF1 and JNK increases expression of stress markers, while blocking the IFN-I signaling increases replication of intracellular bacteria in TNF-primed susceptible macrophages. In contrast, inhibiting transcriptional activity of c-Myc, which is upstream of stress initiation in the sst1S phenotype, improved both macrophage survival and the bacterial control in vitro. We hypothesize that in vivo, Myc downregulation is a part of macrophage reprogramming within inflammatory milieu that allows balancing metabolism and pre-adaption to pathogen encounter.

Despite its focus on effects of a single genetic locus, our study reveals a generalizable paradigm in host – pathogen interactions. Traditionally, susceptibility to a specific pathogen is viewed as an immune defect associated with inability to mount appropriate effector responses. Therefore, studies have been primarily focused on molecules involved in pathogen recognition and immune effector functions. A wealth of information about those molecules and the associated essential mechanisms of host immunity has been obtained studying extreme susceptibility to infections in humans and knockout mice (reviewed in^81,82^. Indeed, avoidance or suppression of host recognition and effector mechanisms by a microbe are strategies essential for establishing infections. Our study demonstrates that subsequent disease progression in immune competent hosts may depend on locally induced macrophage susceptibility that emerge gradually within inflammatory tissue due to an imbalance of macrophage growth, differentiation and stress responses prior to contact with microbes. This explains how successful pathogens may exploit this regulatory failure to bypass mechanisms of resistance in otherwise immune competent hosts. This strategy would ensure survival of both the host and the pathogen and facilitate successful transmission of the later^83^. The origins of aberrant macrophage responses within inflammatory milieu may encompass genetic and non-genetic causes, such as co-infections, metabolic diseases and senescence. Since the sst1-susceptible phenotype in mice closely resembles pathology of human TB, we propose that environmental exposures, metabolic stressors, senescence and co-infections linked to TB risk or severity may also activate this nascent mechanism of infection susceptibility and immunopathology in humans. This concept suggests a novel disease-modifying therapeutic strategy focusing on correcting aberrant response to TNF rather than blocking this essential mediator of host resistance.

## MATERIALS & METHODS

### Reagents

Recombinant mouse TNF was from Peprotech and recombinant mouse IL-3 was from R&D. Mouse monoclonal antibody to mouse TNF (MAb; clone XT22) was from Thermo scientific and isotype control and mouse monoclonal antibody to mouse IFNb (Clone: MAR1-5A3) was from eBiosciences. BAY 11-7082, Phenylbutyrate sodium (PBA), rapamycin were from Enzo Life sciences. SB203580, SP600125 and C16 were obtained from were from Calbiochem. JQ1, Flavopiridol, 10058-F4 were from Tocris. ISRIB, poly I:C, LPS from E. coli(055:B5), Triptolide and BHA were obtained from Sigma. BX-795 was from Invivogen. RHT was kindly provided by Aaron Beeler, BU CMD and Chemistry Department.

### Animals

C57BL/6J and C3HeB/FeJ inbred mice were obtained from the Jackson Laboratory (Bar Harbor, Maine, USA). The C3H.B6-sst1, C3H.scid and C3H.B6-sst1, scid mouse strains were generated in our laboratory as described previously^14,17,84^. The B6.C3H-sst1(B6J.C3-sst1^C3HeB/Fej^Krmn) congenic mice were created by transferring the *sst1* susceptible allele from C3HeB/FeJ mouse strain on the B6 (C57BL/6J) genetic background using twelve backcrosses (referred to as B6-sst1S in the text). All experiments were performed with the full knowledge and approval of the Standing Committee on Animals at Boston University and Johns Hopkins University in accordance with relevant guidelines and regulations.

### BMDMs culture and infection with F. tularensis LVS

Isolation of mouse bone marrow and culture of BMDMs were carried out as previously described^85^. TNF-activated macrophages were obtained by culture of cells for various times with recombinant mouse TNF (10 ng/ml). F. tularensis live vaccine strain (LVS) were grown in Brain-Heart Infusion broth overnight, harvested and then diluted in media without antibiotics to get the desired MOI. BMDM were seeded in tissue culture plates. Cells were treated with TNF and inhibitors were added after 4hrs of TNF addition. After 24hrs of treatment, cells were infected at indicated MOI. The plates were then centrifuged at 500× g for 15 minutes and incubated for 1hr at 37°C with 5% CO2. Cells were then washed with fresh media, and incubated for 45 min at 37°C with media containing gentamicin (50 μg/mL) to kill any extracellular bacteria. Cells were washed again and cultured in presence of inhibitors and TNF in DMEM/F12 containing 10% FBS medium without antibiotic at 37°C in 5% CO2 for 24hrs.

### Bacterial DNA isolation and quantification

For isolation of LVS DNA from 96-well plates, cells were lysed with 50 ul of 250mM NaOH, 0.2 mM EDTA and kept at RT for 10 mins. The samples were then heated for 45 mins at 95^°^C and neutralized with 50 ul 40mM Tris-HCL, pH 8.0. DNA was then isolated by purification with magnetic beads. TaqMan PCR conditions were carried out according to previous studies(ref). All PCR reactions were performed in a final volume of 20ul and contained Taqman Environmental mastermix (Applied Biosystems) at a 1X final concentration, probe (250nM), and primers (450nM). The primers used were FopAF: ATCTAGCAGGTCAAGCAACAGGT, FopAR: GTCAACACTTGCTTGAAC-ATTTCTAGATA, and the probe FOPAP: CAAACTTAAGACCACCACCCACATCCCAA. Thermal cycling conditions were 50°C for 2 min, 95°C for 10 min, 45 cycles at 95°C for 10 s and 60°C for 30 s, and then 45°C for 5 min.

### Animal infections and imaging

Mice were infected by Mtb H37Rv using a Glas-Col chamber, and mice were sacrificed for enumeration of Mtb CFU in lungs and spleen day 1 counts as well as subsequent time points as previously described^86^. H&E stained tissue sections were imaged by high resolution digital microscopy and lesion scoring was performed as described^87^. High resolution PET/CT imaging was conducted using a Mediso nanoScan instrument. Image analysis to quantify disease burden in lungs was performed using previously reported algorithms^88^.

### Immunoblotting

To monitor the Ipr1 protein levels we have developed Ipr1 peptide-specific rabbit polyclonal antibodies, which recognized the Ipr1 protein of predicted length on Western blots (ref Sc Report paper). BMDM’s were subjected to treatments specified in the text. Nuclear extracts were prepared using the nuclear extraction kit from signosis. Whole cell extracts were prepared by lysing the cells in RIPA buffer supplemented with protease inhibitor cocktail and phosphatase inhibitor I and III (Sigma). Equal amounts (30 μg) of protein from whole-cell extracts was separated by SDS-PAGE and transferred to PVDF membrane (Millipore). After blocking with 5% skim milk in TBS-T buffer [20 mMTris–HCl (pH 7.5), 150 mM NaCl, and 0.1% Tween20] for 2 hour, the membranes were incubated with the primary antibody overnight at 4 °C. Bands were detected with enhanced chemiluminescence (ECL) kit (Perkin Elmer). Stripping was performed using WB stripping solution (Thermo scientific). The loading control β-actin (Sigma, 1:2000) was evaluated on the same membrane. The Ipr1 -specific rabbit anti-serum was generated by Covance Research Products, Inc. (Denver, CO, USA)(1:500) as described previously^85^. The Ipr1 monoclonal antibodies were generated using Ipr1 peptides from Abmart. ATF4, ATF3, Gadd34, c-Myc, Daxx, p21, PKR and phospho-PKR antibodies were obtained from Santa Cruz biotechnology:_rabbit polyclonal-anti-p-PKR (sc101783, Santa Cruz), mouse monoclonal-anti-PKR (sc6282, Santa Cruz) were used at dilution factor of 1:150 and 1:200, respectively. IRF1, IRF3 (1:1000), p38, p-p38, JNK, p-JNK antibodies were obtained from Cell signaling. Hspa1a (1:1000) antibody was obtained from R&D. ß-actin (1:2000) was obtained from Sigma. Bim and DHCR7 were obtained from Abcam.

### RNA Isolation and quantitative PCR

Total RNA was isolated using the RNeasy Plus mini kit (Qiagen). cDNA synthesis was performed using the SuperScript II (Invitrogen). Real-time PCR was performed with the GoTaq qPCR Mastermix (Promega) using the CFX-90 real-time PCR System (Bio-Rad). Oligonucleotide primers were designed using Primer 3 software (**Supplementary Table S1**) and specificity was confirmed by melting curve analysis. Thermal cycling parameters involved 40 cycles under the following conditions: 95 °C for 2 mins, 95 °C for 15 s and 60 °C for 30 s. Each sample was set up in triplicate and normalized to RPS17 or 18S expression by the DDCt method.

### Immunofluorescence microscopy

Cells were fixed with 4% paraformaldehyde for 15 min at RT, cells were permeabilised with 0. 25% Triton-X for 30 min and then blocked for 20 min with goat-serum (2.5%). Cells were incubated with primary antibodies [mouse monoclonal antibodies against J2 (1:3000), overnight at 4^°^C in 2.5% goat serum, and incubated with Alexa Fluor 488-conjugated donkey anti-mouse IgG (excitation/emission maxima ~ 490/525 nm) (1:1000, Invitrogen) secondary antibody for 2 hrs. Images were acquired using Leica SP5 confocal microscope. All images were processed using Image J software.

### Hoechst/PI Staining Method for cell cytotoxicity

For cell viability assays BMDM were plated in 96 well tissue culture plates (12000 cells/well) in phenol-red free DMEM/F12 media and subjected to necessary treatments. Hoechst (Invitrogen, 10 μM) and PI (Calbiochem, 2 μM) were added. The plates were kept at 37^°^C for 15 min and read in the celigo cell cytometer. The % of total and dead cells was calculated for each treatment.

### Transcription factor profiling analysis

Each array assay was performed following the procedure described in the TF activation profiling plate array kit user manual (Signosis, Inc, FA-001). 10 ug of nuclear extract was first incubated with the biotin labeled probe mix at room temperature for 30 min. The activated TFs were bound to the corresponding DNA binding probes. After the protein/DNA complexes were isolated from unbound probes, the bound probes were eluted and hybridized with the plate pre-coated with the capture oligos. The captured biotin-labeled probes were then detected with Streptavidin– HRP and subsequently measured with the TECAN microplate reader.

### Gel shift assay

The nuclear extracts with 12 h of TNF treatment in sst1R and sst1S were chosen for gel shift assay analysis with EMSA kits (Signosis Inc). The TF DNA binding probe sequences are listed below.

1. AP1: CGCTTGATGACTCAGCCGGAA
2. c-Myc: AGTTGACCACGTGGTCTGGG

The sequences that we used as probes for gel shift assay are identical to those we used as the probe mix for TF activation profiling array assay. 5ug nuclear extracts were incubated with 1 × binding buffer and biotin-labeled probe for 30 min at room temperature to form protein/DNA complexes. The samples were then electrophoresed on a 6 % polyacrylamide gel in 0.5 % TBE at 120 V for 45 min and then transferred onto a nylon membrane in 0.5 % TBE at 300 mA for 1 h. After transfer and UV cross-linking, the membrane was detected with Streptavidin–HRP. The image was acquired using a FluorChem imager (Alpha Innotech Corp).

### siRNA knockdown

Gene knockdown was done using GenMute (SignaGen) and Flexitube Genesolution siRNAs from Qiagen. All star negative control siRNA (SI03650318) from Qiagen was used as a negative control. sst1S and sst1R BMDMs were seeded into 6-well plates at a density of 2.5 × 10^5^ per well and grown as mentioned before. Shortly before transfection, the culture medium was removed and replaced with 1 ml complete medium, and the cells were returned to normal growth conditions. To create transfection complexes, 15 nM siRNA (pool of 4 siRNAs) in 1 × GenMute buffer (total 500 ml) was incubated with 1.5 μl of GeneMute transfection reagent for 15-20 minutes at room temperature. The complexes were added drop-wise onto the cells. Cells were incubated with the transfection complexes for 24 hours at 37^°^ in 5% CO2. After 24hrs cells were washed to remove siRNA and replaced with fresh media. TNF (10ng/ml) was added for 24hrs and BMDMs were harvested as outlined below. siRNA pools included: Irf1 (GS16362), Irf3 (GS54131), Irf7(GS54123)

### ELISA

Supernatants were collected from mouse macrophages after 24hrs of stimulation with TNFa or poly IC. IFNβ was measured using the mouse IFNβ ELISA kit from PBL Assay Science. ELISAs were done as recommended by the manufacturer.

## Acknowledgement

This work was sponsored by R01 HL133190 and R01 HL126066 to IK and WB. The authors are grateful to Drs. Robert Silverman, Karen Elkins and Benjamin Wolozin for helpful discussions, and to Adam Gover and Somak Ray for the analyses of microarray and proteomics data, respectively, and to Dr. Kathleen Mulka for assistance with histologic scoring and quantification.

The authors declare no competing interests.

