## Supplementary figures and images for "The integrated stress response mediates type I interferon driven necrosis in *Mycobacterium tuberculosis* granulomas"

### Supplemental Figure 1

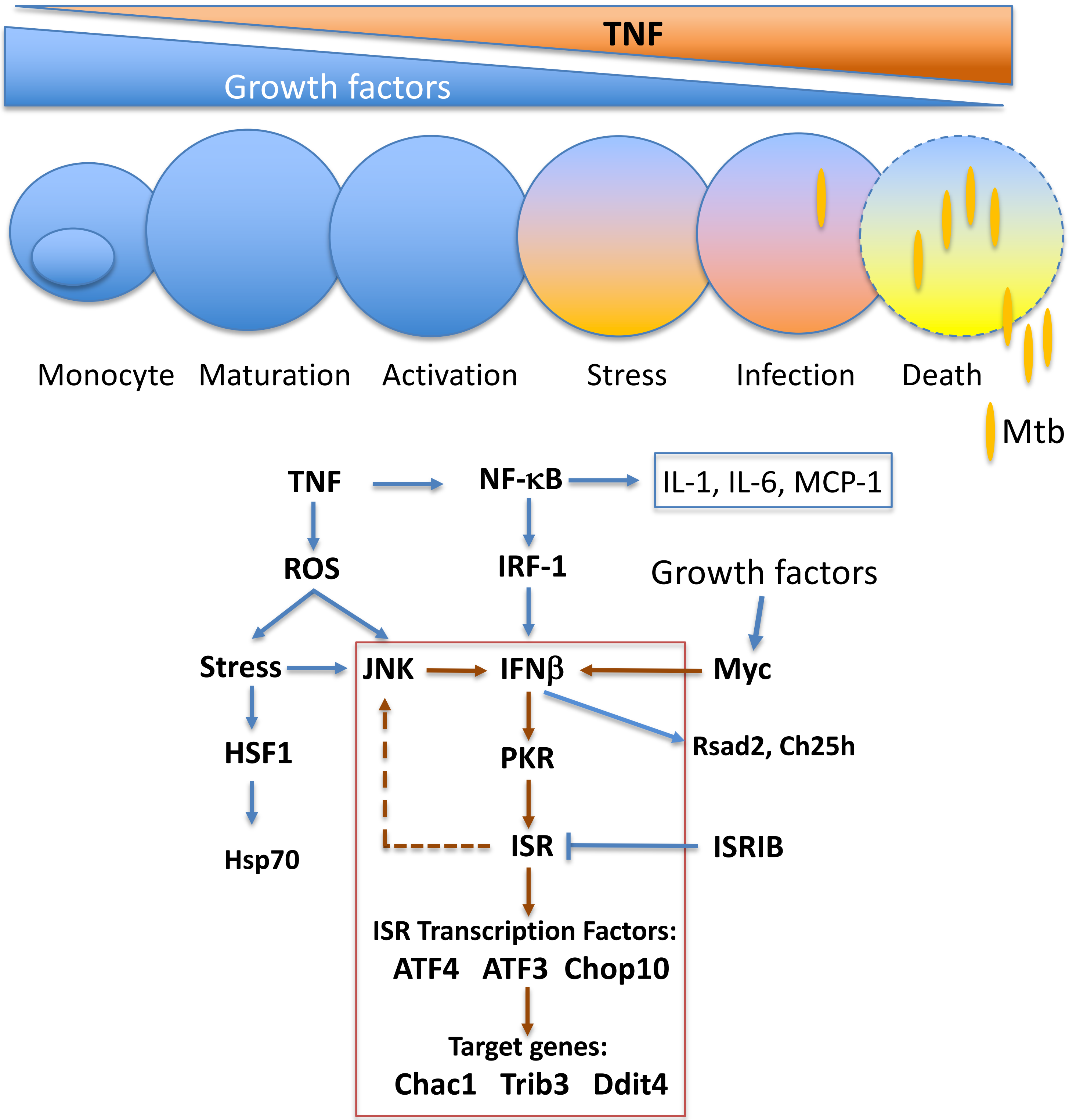

### Supplemental Figure 2

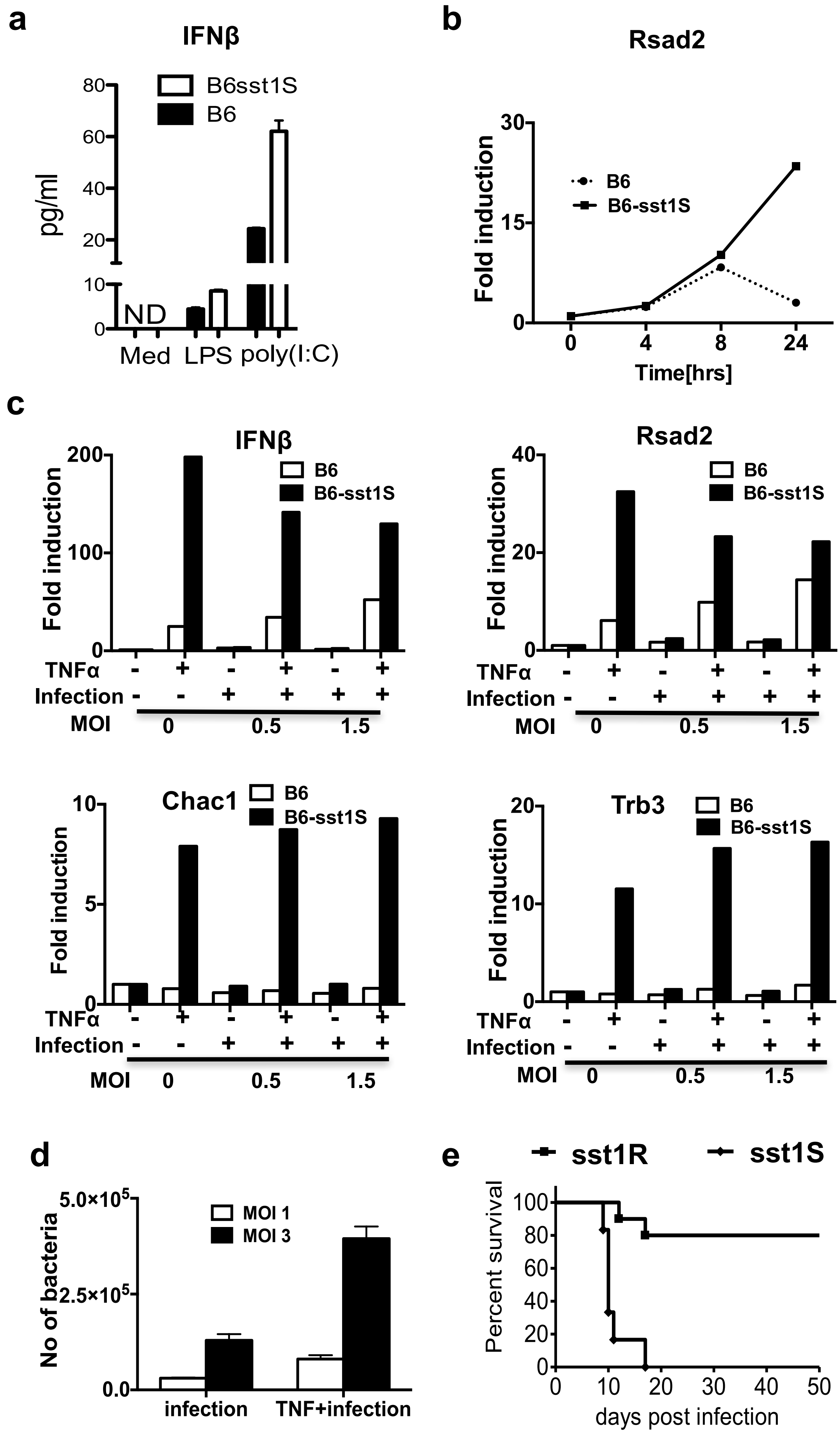

### Supplemental Figure 3

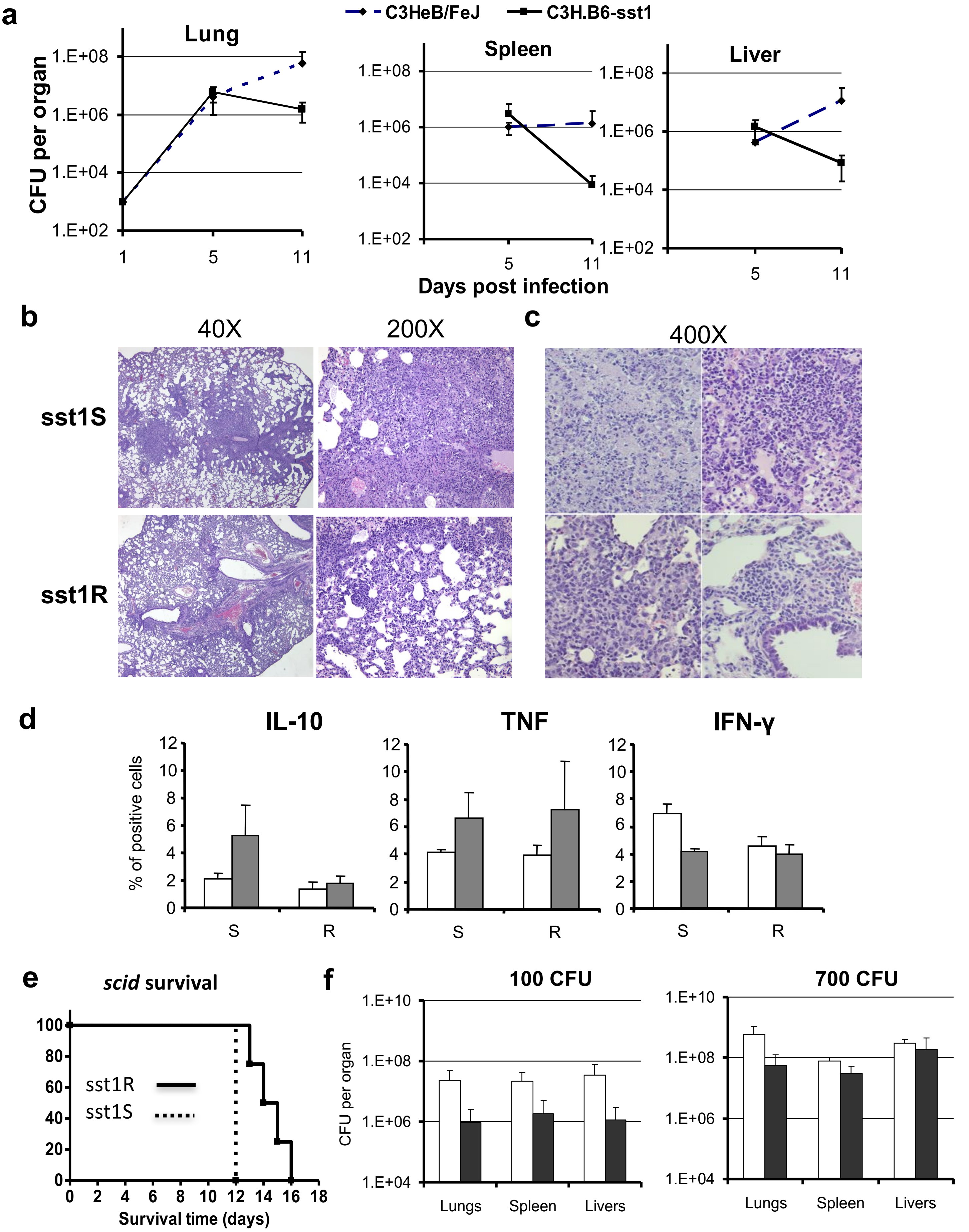

### Supplemental Figure 4

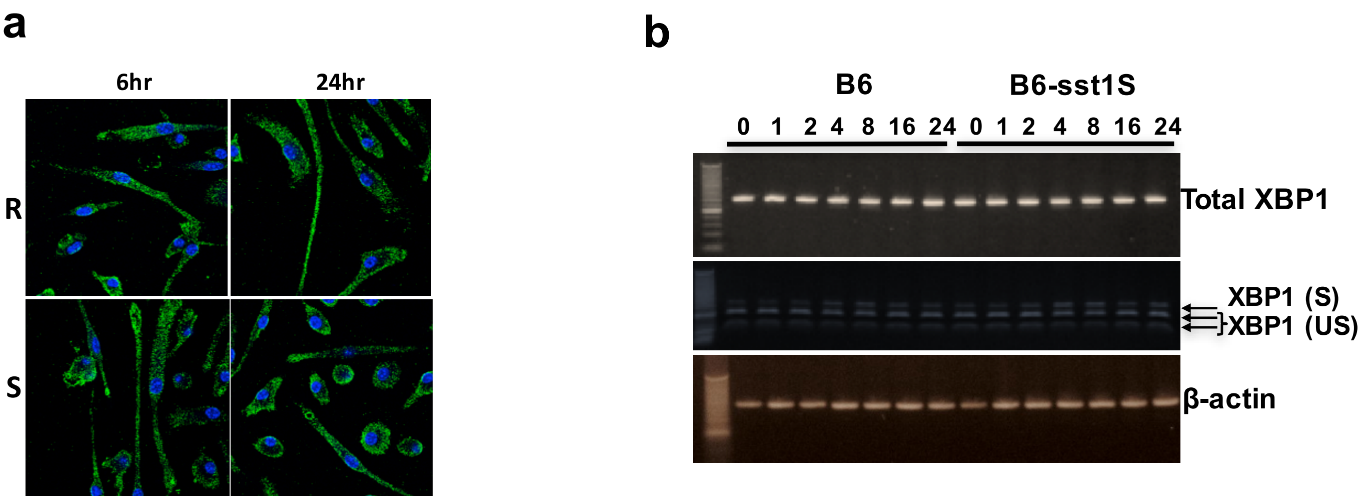

### Supplemental Figure 5

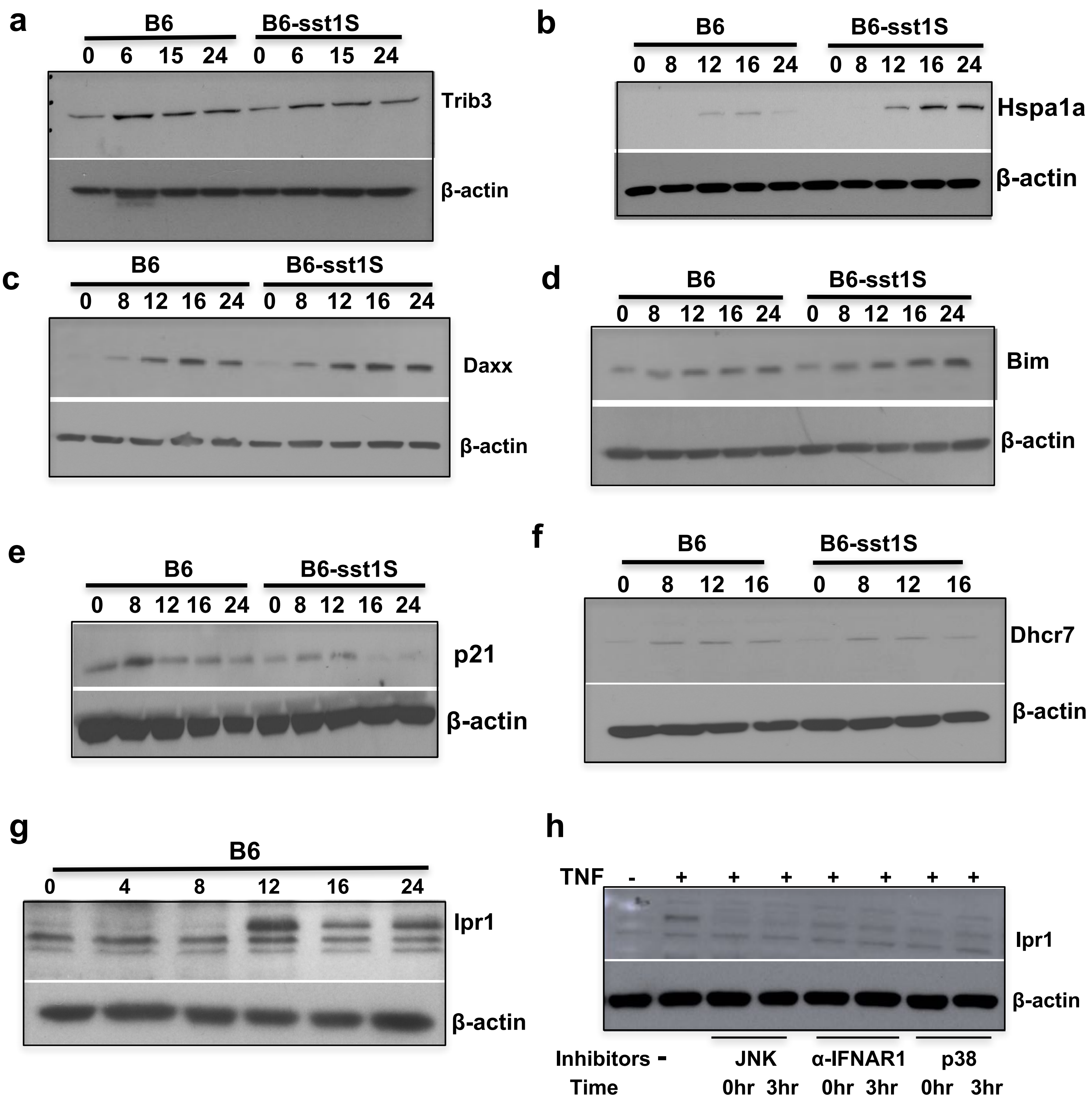

### Supplemental Figure 6

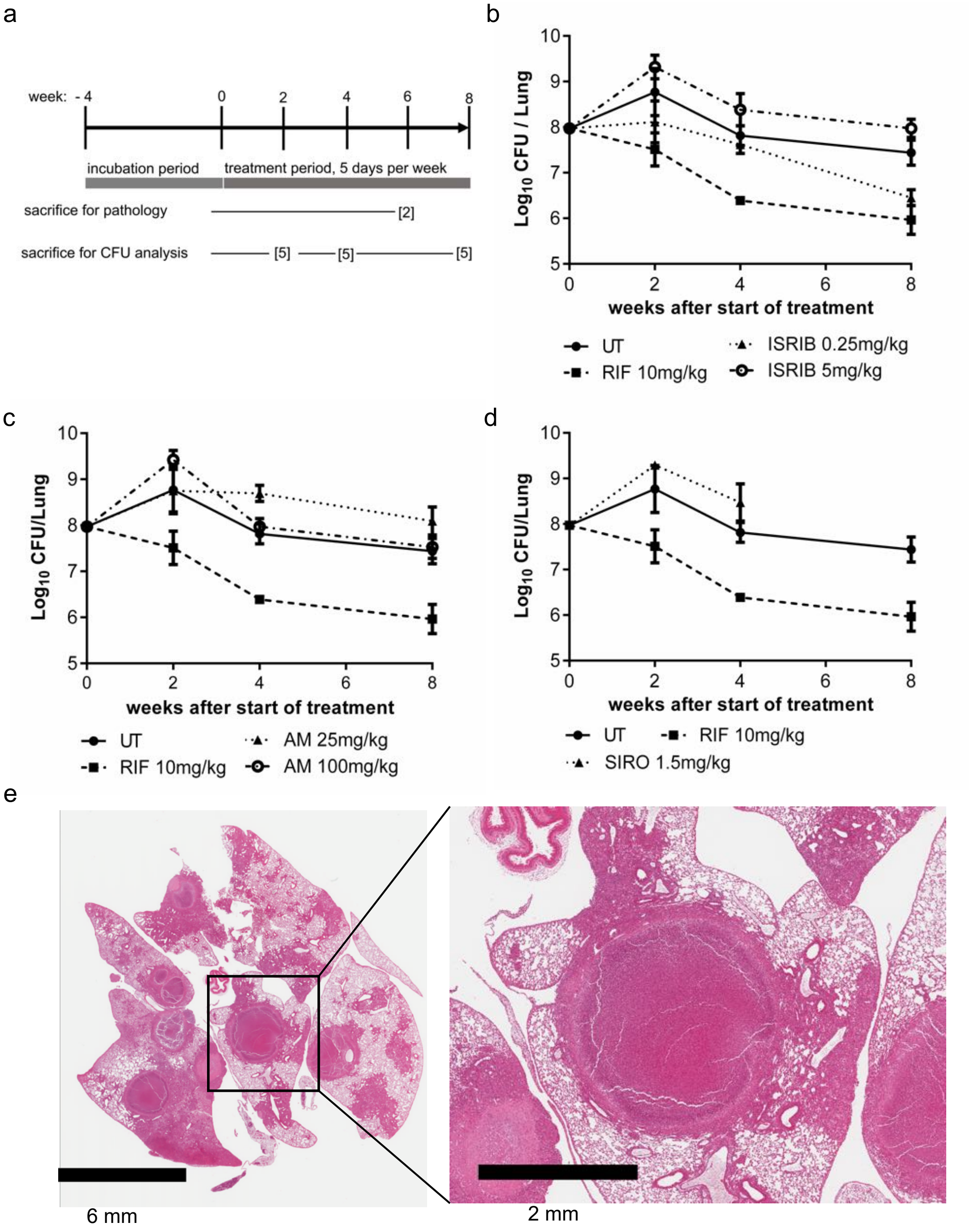

### Supplemental Figure 7

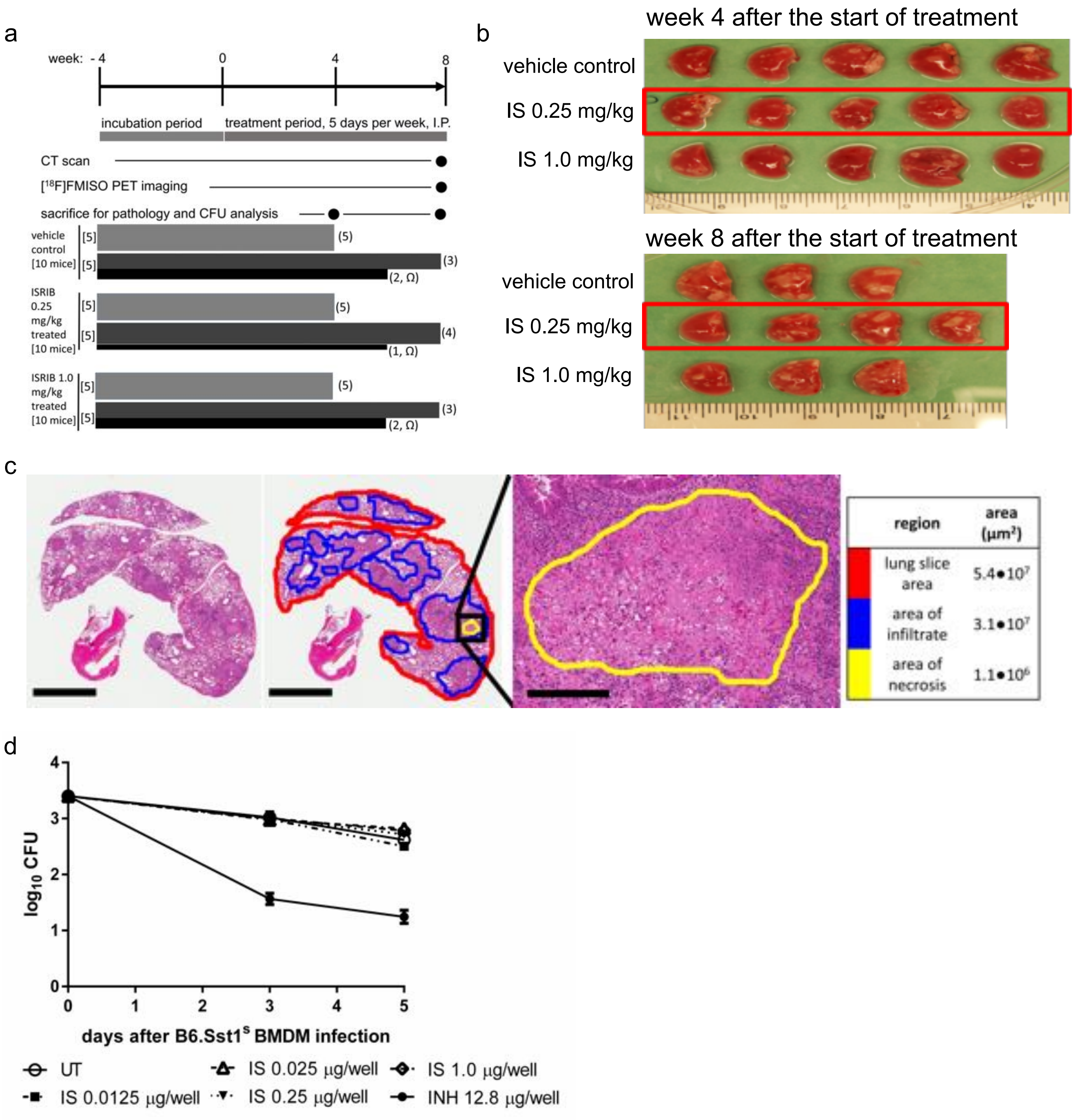

### Supplemental Figure 8

a

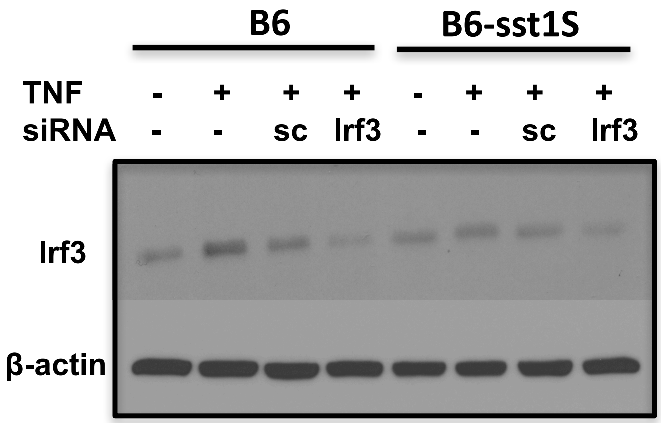


b

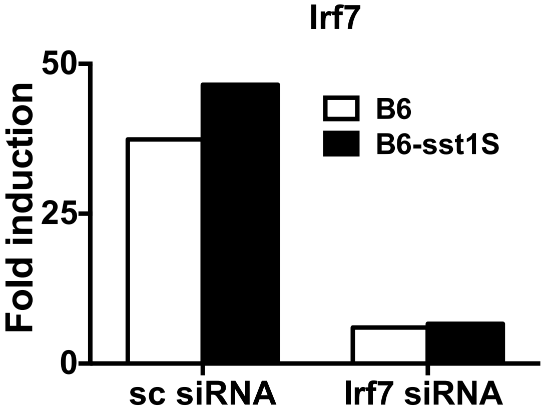


Suppl.Fig.8

### Supplemental Figure 9

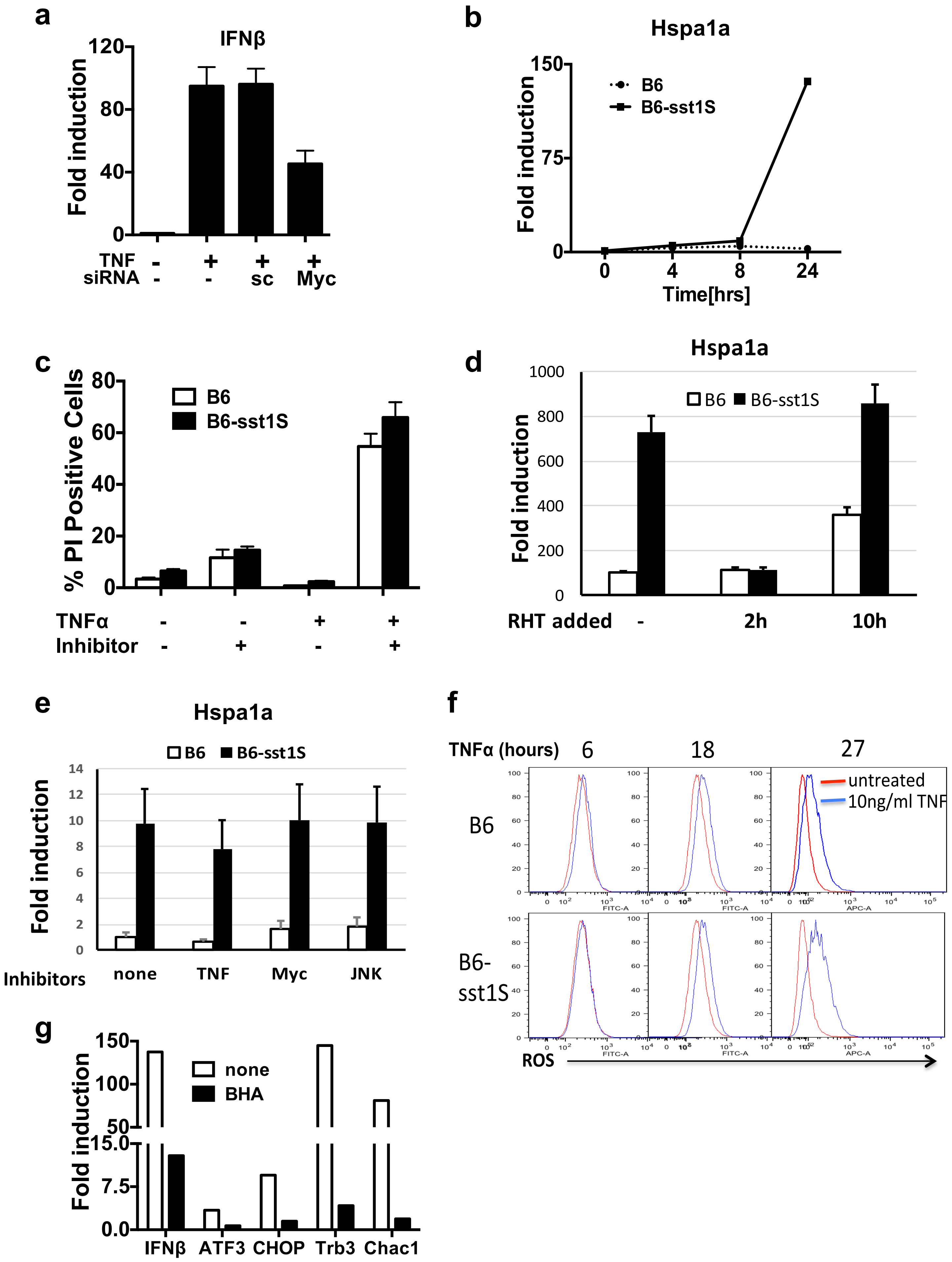
