## Supplemental Figure Legends for "The integrated stress response mediates type I interferon driven necrosis in *Mycobacterium tuberculosis* granulomas"

**Supplementary Figure 1.** Aberrant activation of the Integrated Stress Response (ISR) by TNF in sst1S macrophages via stress-dependent super-induction of type I IFN pathway.

Blue lines – canonical TNF-activated pathways

Orange lines – mechanisms of IFNβ pathway super-induction in B6-sst1S macrophages

Orange box – PKR-mediated integrated stress response pathway activation in B6-sst1S macrophages

**Supplementary Figure 2.**

**a)** IFNβ concentrations in supernatants of B6wt and B6-sst1S BMDM treated with 100ng/ml LPS and 1ug/ml poly IC for 24h, as determined by ELISA.

**b**) Time course of Rsad2 mRNA expression in B6wt and B6-sst1S BMDM treated with 10ng/ml TNF (representative of two independent experiments).

**c**) Comparison of the effects of TNF and *F.t.*LVS infection on IFN-I and stress response gene expression in B6-sst1S BMDM. Macrophages, either naïve or primed with 10ng/mL of TNFα for 16 h, were infected with *F.t.* LVS at MOI 0.5 and 1.5 for 24hrs. IFNβ, Rsad2, Trb3 and Chac1 mRNA expression was measured using qRT-PCR. Real time PCR data is normalized to expression of 18S rRNA and presented relative to expression in untreated cells (set as 1).

**d**) Effect of TNF priming on *F.t.* LVS control by the B6-sst1S BMDM. The macrophages, either naïve or primed with 10ng/mL of TNFα for 16hrs, were infected with *F.t.* LVS at MOI 1 and 3 for 24 h. The bacterial loads were determined using qPCR.

**e**) Survival of the $sst1^R$ and $sst1^S$ inbred mouse strains after aerosol infection with 1,600 CFU of *F.t.* LVS. Six mice per group were used in each strain.

**Supplementary Figure 3.**

**a**)  Kinetics of *F.t.* LVS growth in the lungs, spleens and livers of the *sst1*$^{R}$ and *sst1*$^{S}$ mice after the aerosol infection with 1,600 CFU of F. LVS. Four mice per group were sacrificed at each time point for CFU determination using plating of serial dilutions of lung homogenates.

**b**) Histopathology of the lungs of *sst1*$^{S}$ (upper panels) and *sst1*$^{R}$ (lower panels) 11 days post aerosol infection with 1,600 CFU of F. LVS. H&E staining, magnification X40 (left panels) and X200 (right panels).

**c**) Histopathology of the lungs of *sst1*$^{S}$ (upper panels) and *sst1*$^{R}$ (lower panels) 11(left panels) and 16 (right panels) days post aerosol infection with 1,600 CFU of F. LVS. H&E staining, original magnification - X400.

**d)** Intracellular cytokine staining of the lung cells isolated 5 (white bars) or 10 (grey bars) days post aerosol infection with 300 CFU of *F.t.* LVS. Three mice per group were used in the experiment. S - sst1$^{S}$; R - sst1$^{R}$. The data is representative of two independent experiments.

**e)** Survival of the *sst1* congenic *scid* mice after aerosol infection with 700 CFU of *F.t.* LVS by aerosol. The C3H.B6-*sst1,scid* (*sst1*$^{R}$ - solid line) and C3H.*scid* (*sst1*$^{S}$ - dashed line) mouse strains were used in this experiment. Six mice per group were used for each strain.

**f**) *F.t.* LVS burdens in the organs of the *sst1*$^{R}$ and *sst1*$^{S}$ mice (C3H.B6-*sst1,scid* and C3H.*scid*, respectively) 11 days after aerosol infection with 100 or 700 CFU of F. LVS. Four mice per group were used for each data point.

**Supplementary Figure 4.**

**a**)  Staining with dsRNA-specific J2 antibodies of B6wt and B6-sst1S BMDM treated with 10ng/ml TNF for 6 and 24 h. Cells were stained with J2 antibody (green); nuclei are counterstained with DAPI (blue). All microscopic images represent data from two independent experiments.

**b**) Time course analysis of XBP-1 mRNA expression and splicing in TNF-stimulated B6wt and B6-sst1S BMDM. Total RNA from TNF stimulated B6wt and B6-sst1S BMDM were amplified using RT-PCR and digested with Pst1as described in Methods. The resultant PCR products were separated by electrophoresis on 2% agarose gel. The PCR products of spliced XBP1(S) remained intact, whereas the PCR products of unsplicedXBP1(U) mRNA were cut into two fragments as indicated by arrows.

**Supplementary Figure 5.** Validation of proteomics data using immunoblot.
The kinetics of proteins differentially expressed in B6-sst1S and B6wt macrophages after stimulation with10ng/mL of TNF: Trib3(**a**), Hspa1a (**b**), Daxx(**c**), Bim(**d**), p21(**e**) and DHCR7(**f**). Whole cell extracts isolated at indicated times were used for the Western blots. **g**) The kinetics of the IPR1 protein expression in B6wt macrophages after TNF stimulation. **h**) Effects of JNK and p38 inhibitors and IFNAR blockade on IPR1 protein induction in B6wt BMDM treated with 10ng/mL of TNF$\alpha$ for 24 h. The inhibitors were added at the beginning (0hr) or after 3hrs of TNF stimulation. Immunoblotting was carried out using Ipr1 polyclonal antibody and represents two independent experiments.

**Supplementary Figure 6.** Effect of small molecule inhibitors on TB progression in C3HeB/FeJ mice.
**a**) Overview of experiments and data presented in b – e. [#] indicates number of mice/analysis group at each endpoint.
(**b - d**) *M. tb*-infected C3HeB/FeJ mice were treated with ISRIB (IS) at doses 0.25 or 5 mg/kg, Amlexanox (AM) at doses 25 or 100 mg/kg, sirolimus (SIRO) at dose 1.5 mg/kg, and untreated (UT). All doses were given 5 days per week during the treatment period. Data plotted as means ± SEM.

**e**) Histopathology exemplifying necrotic lung granulomas in a mouse treated with Amlexanox 25mg/kg at week 6 of treatment (H&E staining).

**Supplementary Figure 7.** Effect of ISRIB on TB progression in B6-sst1S mice and on *M.tb* replication in B6-sst1S BMDMs.

**a**) Experimental overview of mouse chronic infection model used to test effects of ISRIB monotherapy on *M. tb* bacterial growth. At left, [#] indicates number of mice in each experimental group and subgroup in the experimental plan. At right, (#) indicates number of mice in each analysis group. (#, Ω) indicates number of mice excluded from analysis because of morbidity and mortality based on IRB and study criteria.

**b**) Lung gross pathology of B6-sst1S mice infected with *M. tb* following 4 and 8 weeks of ISRIB treatment (8 and 12 weeks post infection).

**c**) Overview of quantitative lung pathology analysis. Areas were drawn by eye and validated by a veterinary pathologist. Analysis was conducted using Aperio ImageScope (Leica Biosystems)

**d**) BMDM infection ISRIB dose ranging experiment. B6-sst1S mice bone marrow derived macrophages were infected with *M. tb* and treated with ISRIB (0.0125, 0.025, 0.25 or 1 μg/ml) 1 hour after infection. Bacillary loads were enumerated in day 0, 3 and 5 after infection. INH=isoniazid, UT=untreated infected macrophage. IS = ISRIB. Data plotted as means±SD.

**Supplementary Figure 8.**

**a**) IRF3 expression and validation of IRF3 knockdown using immunoblotting with IRF3-specific antibodies (sc – scrambled control; Irf3 – specific siRNA).

**b**) Validation of Irf7 knockdown in B6wt and B6-sst1S BMDM. Irf7 mRNA expression was quantified using qRT-PCR, normalized to expression of 18S rRNA and presented relative to expression in untreated cells (set as 1). The data are representative of two independent experiments.

**Supplementary Figure 9.**

**a**) Effect c-Myc knockdown on IFNβ mRNA expression in B6-sst1S BMDM stimulated with TNF (10 ng/ml) for 18 h.

**b**) Timecourse of Hspa1a mRNA expression in B6 and B6-sst1S BMDM stimulated with TNF (10ng/ml).

**c**) Synergistic effect of TNF and HSF1 inhibitor KRIB11 on macrophage death. B6 and B6-sst1S BMDM were treated with 10ng/ml TNFα for 24 hrs in presence or absence of 10uM KRIB11. Cell death was assessed using automated microscopy as % of PI positive cells. Data represents results from two independent experiments.

**d**) Effect of the HSF1 and translation inhibitor rohitinib (RHT) on Hspa1a mRNA expression in B6wt and B6-sst1S BMDM treated with 10ng/mL of TNF for 24 hrs. RHT (300uM) was added at 2 and 10 h of TNF stimulation. Gene expression data are normalized to expression of 18S mRNA and presented relative to expression in untreated cells (set as 1). Results represent data from three independent experiments.

**e**) No effects of TNF blockade, c-Myc or JNK inhibitors added 12h of TNF stimulation on Hspa1a mRNA expression at 16 h in the B6-sst1S BMDMs.

**f)** ROS production in B6wt and B6-sst1S BMDM stimulated with 10ng/ml TNFα for the indicated times. FACS analysis was performed using DCFDA (dichlorofluorescindiacete) probe (Abcam).

**g)** Effect of ROS inhibition with BHA on IFNβ and ISR gene expression in B6-sst1S BMDM treated with 10ng/mL of TNF for 24 h. The gene expression is normalized to expression of 18S rRNA and presented relative to expression in untreated cells (set as 1).
